# From *in silico* to *in vitro*: Wŭhàn sharpbelly bornavirus infects and persists in cyprinoform cells

**DOI:** 10.1101/2025.03.26.645528

**Authors:** Mirette I. Y. Eshak, Angele Breithaupt, Birke A. Tews, Christine Luttermann, Kati Franzke, Marion Scheibe, Sören Woelke, Martin Beer, Dennis Rubbenstroth, Florian Pfaff

**Author notes:** Corresponding author: Florian Pfaff.

## Abstract

Our recent study using *in silico* data mining identified novel culterviruses (family: *Bornaviridae*) in fish, including a variant of Wŭhàn sharpbelly bornavirus (WhSBV) in grass carp kidney and liver cell lines. Here, metagenomic sequencing of different fish cell lines revealed WhSBV in two cell lines from grass carp (*Ctenopharyngodon idella;* order: Cypriniformes). Using these cell lines, we investigated the ability of WhSBV to infect and establish persistent infection in other cell lines from bony fish (Cypriniformes, Chichliformes, Salmoniformes, Centrarchiformes and Spariformes), reptiles (Testudines and Squamata), birds (Galliformes) and mammals (Primates and Rodentia). WhSBV showed efficient replication and a time-dependent increase in viral RNA levels in cypriniform cells, whereas replication was limited, confined to single cells, and lacked a clear time-dependent increase in cells from other bony fish and reptiles. No replication was detected in avian and mammalian cells. *In situ* hybridisation and electron microscopy confirmed the presence of viral RNA and particles in infected cypriniform cells. Transcriptomic sequencing revealed minimal innate immune activation during early stages of infection and antiviral response only at later stages, suggesting that WhSBV establishes persistence by evading early immune recognition. In addition, we identified polycistronic viral mRNAs regulated by specific transcriptional start and termination sites and RNA splicing. Viral proteins were detected, confirming previous *in silico* predictions. These findings provide insights into the potential infectivity, persistence mechanisms and transcriptional strategies of WhSBV. This study validates previous findings from *in silico* data mining, further reinforcing its effectiveness as a powerful tool for discovering hidden viruses.

**IMPORTANCE:** Understanding the diversity and host range of viruses is crucial for assessing their ecological role, associated diseases and zoonotic potential. However, many newly discovered viruses are characterised using sequence data alone because isolates are often difficult to obtain. Using cell culture models, this study characterises Wŭhàn sharpbelly bornavirus (WhSBV), a member of the genus *Cultervirus*. Here we demonstrate its ability to establish persistent infection in cypriniform fish cell lines, while exhibiting restricted replication in certain non-cypriniform fish. The identification of polycistronic transcription, splicing events and immune evasion mechanisms advances our understanding of the molecular biology of WhSBV and culterviruses in general. By validating *in silico* predictions, this study highlights the power of computational approaches in uncovering viral diversity. As cypriniform fish include economically important species such as carp, understanding the dynamics of WhSBV host range and infection biology may be crucial for future aquaculture health management.

## INTRODUCTION

The family *Bornaviridae*, within the order *Mononegavirales*, contains four genera: *Orthobornavirus*, *Carbovirus*, *Cultervirus*, and *Cartilovirus* [1–3]. The genus *Orthobornavirus* has a remarkably broad host range, infecting avian, reptilian, and mammalian species [3]. Furthermore, orthobornaviruses include zoonotic viruses, such as Borna disease virus 1 (BoDV-1) and variegated squirrel bornavirus 1, both of which can cause fatal encephalitis in humans [4, 5]. Carboviruses and culterviruses, on the other hand, have only been found in reptiles and fish, respectively, and no zoonotic spill-over has been reported [3, 6, 2, 7]. The genus *Cartilovirus* currently only includes the little skate bornavirus (LSBV), which was discovered using public datasets from the Sequence Read Archive (SRA) [2].

Bornavirids possess enveloped virions that enclose a linear, negative-sense, monopartite RNA molecule of approximately 9,000 nucleotides (nt) [8, 3]. Their genomes encode for at least six open reading frames (ORFs) arranged in the order 3’-N-X/P-M-G-L-5’ for orthobornaviruses or 3’-N-X/P-G-M-L-5’ for carboviruses and culterviruses. These ORFs encode for six viral proteins; nucleoprotein (N), accessory protein (X), phosphoprotein (P), matrix protein (M), glycoprotein (G), and the large protein (L) which contains a RNA-directed RNA polymerase [8, 3]. Recently, the largest bornavirid genome, comprising 11,090 nt, was identified for the cartilovirus LSBV. The LSBV genome encodes two additional ORFs, potentially encoding viral proteins 1 and 2 (vp1 and −2), and represents a unique genomic architecture 3’-N-vp1-vp2-X/P-G-M-L-5’ [2]. Studies of BoDV-1 have demonstrated that its RNA and genome are transcribed and replicated in the host cell nucleus, respectively [9]. In bornavirids, the generation of a diverse array of mRNAs is regulated by differential use of transcription start and termination sites [10], as well as alternative splicing of polycistronic primary transcripts to increase and control their transcriptional complexity [11, 12].

Wŭhàn sharpbelly bornavirus (WhSBV, NC_055169; species *Cultervirus hemicultri*), a member of the genus *Cultervirus*, was initially identified by RNA sequencing of the gut, liver, and gill tissues from a freshwater sharpbelly (*Hemiculter leucisculus* [Basilewsky, 1855], family Xenocyprididae, order Cypriniformes) from China [6, 3]. Our recent study using *in silico* data mining of publicly available sequencing datasets led to the identification of novel fish culterviruses, including the first complete genome of the Murray-Darling carp bornavirus (MDCBV, BK063521) from a goldfish (*Carassius auratus* [Linnaeus, 1758], family Cyprinidae, order Cypriniformes) tissue pool data set and a WhSBV variant (BK063520) detected in grass carp (*Ctenopharyngodon idella* [Valenciennes, 1844], family Xenocyprididae, order Cypriniformes) CIK (kidney) and L8824 (liver) cell lines [2]. The two viruses belong to the same bornavirid species (*Cultervirus hemicultri*) and share 78% nt overall identity. However, current knowledge of these culterviruses remains limited to *in silico* genetic discoveries and bioinformatic predictions, with no virus isolates available for experimental research.

To deepen our understanding of the biology and abundance of culterviruses, we screened 48 fish-derived cell lines for the presence of known or novel culterviruses using metagenomic RNA sequencing. This revealed the presence of WhSBV in two grass carp swim bladder cell lines, with genomes nearly identical to the aforementioned WhSBV variant BK063520 from the liver (L8824) and kidney (CIK) cell lines. This study represents the first investigation into the biology of WhSBV and lays the groundwork for future research of culterviruses.

## MATERIALS AND METHODS

### Nucleic acids extraction and metagenomic sequencing

A total of 48 fish cell lines were provided by the Collection of Cell Lines in Veterinary Medicine (CCLV) at the Friedrich-Loeffler-Institut, Riems, Germany (see Supplementary Table S1 for a complete list of cell lines). Fresh-frozen cell lines were lysed in 1 ml TRIzol LS reagent (Life Technologies) and shaken vigorously for 10 min at room temperature. After addition of 200 µl chloroform (Carl Roth) and centrifuging at 13,000 × g, 10 min, 4°C, 200 µl of the upper aqueous phase was used for RNA extraction. Total RNA was extracted using the Agencourt RNAdvance Tissue Kit (Beckman Coulter) with a KingFisher Flex Purification System (Thermo Fisher Scientific) according to the manufacturer’s instructions. DNase I digestion (Qiagen) was carried out prior to the final elution to remove genomic DNA residues. The quantity of total RNA was determined using the NanoDrop Lite Spectrophotometer (Thermo Fisher Scientific). Poly(A) RNA was isolated using the Dynabeads mRNA DIRECT Purification Kit (Invitrogen) following the manufacturer’s instructions. Fragmentation and strand-specific construction of whole transcriptome libraries was performed using the Collibri Stranded RNA Library Prep Kit for Illumina Systems (Invitrogen). The quality and integrity of the RNA and the final library were verified using the Agilent TapeStation 4150 (Agilent Technologies) with appropriate chips and reagents. The final libraries were quantified using the Qubit dsDNA HS Assay Kit (Invitrogen) in combination with the Qubit 2.0 Fluorometer (Invitrogen). Equimolar amounts of the library were then sequenced using Novaseq 6000 with SP flow cell in single-end 101 bp mode (CeGat GmbH) according to the manufacturer’s instructions.

### Metagenomic analysis

The raw reads obtained from the 48 fish cell lines were trimmed using *TrimGalore!* (v0.6.10; [13]) and *cutadapt* (v4.0; [14]). The metagenomic pipeline *SqueezeMeta* (v1.6.4; [15]) was then used to individually assemble the trimmed reads from each dataset and then sequentially merge them into a single transcriptome (“seqmerge” mode). Poly(A/T) sequences at the sequence termini of the assembled and merged transcripts were then removed using *cutadapt* (v4.0; [14]). The polished transcriptome was then used as input for taxonomic assignment and abundance estimation using *SqueezeMeta* (v1.6.4; [15], database version Sep-3-2023). The results were further analysed using the R (v4.3.1; [16]) package “SQMtools” (v1.6.3; [15]).

### Validation of WhSBV genome termini

Persistently WhSBV-infected grass carp swim bladder (GCSB) cell lines GCSB1441 and GCSB1542 (CCLV no. 1441 and 1542, respectively; see Supplementary Table S1) were grown in non-vented T25 tissue culture flasks (Corning) in a suitable growth medium supplemented with 10% fetal bovine serum (FBS) to 95-100% confluence. Total RNA was extracted using the TRIzol-chloroform method, followed by purification using the RNeasy Mini Kit (Qiagen) and DNase I digestion prior to elution. The quality and quantity of the RNA were assessed using the Agilent TapeStation 4150 in combination with appropriate chips and reagents, and the NanoDrop Lite spectrophotometer, respectively.

The 5’-Rapid Amplification of cDNA Ends (RACE) kit, version 2.0 (Invitrogen), was used to determine the 3’ end of the WhSBV genome. Reverse transcription (RT) was performed on the extracted RNA using a gene-specific primer WhSBV-945-R (0.4 μM) (see Supplementary Table S2 for a complete list of primers). The resulting cDNA was then digested with RNase cocktail (Invitrogen) at 37°C for 20 min and subsequently purified using AMPure XP beads (Beckman Coulter). A tailing reaction was then performed using dCTP and dATP (New England Biolabs). A second round of amplification was performed using gene-specific primers WhSBV-777-R (0.2 μM) and the AAP primer (0.2 μM) for purified C-tailed cDNA or the adapter primer (AP) (0.2 μM) for purified A-tailed cDNA (see Supplementary Table S2). The hemi-nested PCR was performed with gene-specific primers WhSBV-545-R (0.2 μM) and the AUAP primer (0.2 μM) (see Supplementary Table S2). To identify the 5’ end, total RNA samples were treated with the *E. coli* Poly(A) Polymerase Kit (New England Biolabs) according to the manufacturer’s instructions in order to add poly(A) tails. The poly(A) tailed RNA were then purified using AMPure XP beads. This purified poly(A) tailed RNA was used as input to the 3’-RACE system for rapid amplification of cDNA ends (Invitrogen), using the AP primer (see Supplementary Table S2) for RT. The cDNA was then digested with an RNase cocktail and purified using AMPure XP beads. For the second round of amplification, six different gene-specific primers (WhSBV_7388F, WhSBV_7528F, WhSBV_7911F, WhSBV_8215F, WhSBV_8737F, WhSBV_8901F; 0.2 μm) were combined with the AUAP primer (see Supplementary Table S2), for individual reactions. The resulting PCR amplicons were separated by 2% agarose gel electrophoresis. The remaining product was purified with AMPure XP beads and sequenced using the BigDye Terminator v1.1 Cycle Sequencing Kit on a 3500 Genetic Analyzer (Applied Biosystems). The final WhSBV genome sequences were uploaded to Genbank under accession number PV171101.

### RT-qPCR and qPCR assays for specific detection of WhSBV

For the following assays, persistently WhSBV-infected GCSB1441 and GCSB1542 cells and the WhSBV-uninfected GK (0119) cell line (see Supplementary Table S1) were grown to 95-100% confluence in non-vented T25 tissue culture flasks using appropriate growth medium supplemented with 10% FBS. Cell pellets were harvested using Alsever’s trypsin-versus-versus-solution (ATV; FLI, Riems) after discarding the supernatant, and RNA/DNA was extracted using the NucleoMag VET kit (Macherey-Nagel) according to the manufacturer’s instructions.

WhSBV-specific reverse transcription quantitative polymerase chain reaction (RT-qPCR) assays were established by designing primers and probe sets targeting the viral N, P, G, M and L genes (see Supplementary Table S2, for a complete list of primers and probes). WhSBV-specific RNA (and potentially DNA) was detected using the AgPath-ID one-step RT-PCR reagents (Thermo Fisher Scientific). Briefly, 5 µl of extracted RNA was reverse transcribed and amplified in a reaction mix of 25 µl total volume containing the specific primer/probe mix (final primer and probe concertation: 0.4 and 0.2 µM, respectively). For the internal control, primers ACT-1030-F (final concentration: 0.2 µM) and ACT-1135-R (0.2 µM) along with probe ACT-1081-HEX (0.2 µM) were included in each reaction to target the cellular actin beta (ACTB) gene [17]. The reaction was performed on a Bio-Rad CFX96 qPCR cycler (Bio-Rad) using the following cycling conditions: 45°C for 10 min, 95°C for 10 min, 42 cycles of 95°C for 15 s, 57°C for 20 s and 72°C for 30 s. The efficiency and limit of detection of the WhSBV G gene RT-qPCR assay was tested using serial dilutions. A Cq cut-off value of 37 was set as limit of detection for all performed assays.

To exclude the presence of WhSBV sequences as endogenous viral elements (EVE) integrated into the cellular genome of GCSB1441, GCSB1542, and GK, we specifically tested extracted DNA from these cells using a non-reverse transcription qPCR system. WhSBV cDNA from the 3’-RACE experiment, as previously described, was used as a positive control. The Maxima Probe/ROX qPCR Master Mix (Thermo Fisher Scientific) was used according to the manufacturer’s protocol, targeting the viral genes N, P, G, M and L (final primer and probe concertation: 0.4 and 0.2 µM, respectively) and the cellular *ACTB* gene (final primer and probe concertation: 0.2 and 0.2 µM, respectively). The reaction was performed on a Bio-Rad CFX96 qPCR cycler using the following cycling conditions: 50°C for 2 min, 95°C for 10 min, 40 cycles of 95°C for 15 s, 60°C for 30 s and 72°C for 30 s.

### Optimisation of WhSBV inoculum preparation

Five different methods were evaluated to prepare the WhSBV inoculum in order to determine the most efficient and reliable protocol for subsequent experiments. Persistently WhSBV-infected GCSB1441 cells were maintained in non-vented T75 (75 cm²) culture flasks (Corning) at 26°C until reaching 95-100% confluency in growth medium supplemented with 10% FBS. For “Method 1”, 5 ml of supernatant (equivalent to 1 ml per 15 cm² of infected cell layer) were retained, and the cells were subjected to three freeze/thaw cycles freezing at −80°C and thawing at room temperature for 30 minutes each. For “Method 2”, after three freeze/thaw cycles as described above, the resulting cell suspension was collected in a Falcon tube (Sarstedt), sonicated (Branson Sonifier 450), and centrifuged at 1,200 × *g* for 20 minutes at 4°C, and only the supernatant was collected. For “Method 3”, cell lines were suspended in 5 ml supernatant, which was subsequently sonicated and centrifuged as described above, and only the supernatant was collected. For “Method 4”, the culture medium was replaced with 5 ml hypertonic medium containing 150 mM NaCl. After, 2 h of incubation at 26°C, the cells were scraped using a cell scraper, and the cell suspension was centrifuged as previously described and only the supernatant was collected. For “Method 3” 5 ml of untreated supernatant was collected. All inocula were stored at −80°C until further use.

In order to test the infectivity of the different inocula, CCB cells (see Supplementary Table S3) were seeded in 12-well plates (3.8 cm^2^ cell layer per well; Corning) and grown to 95-100% confluency in the appropriate medium supplemented with 10% FBS at 26°C. Inoculation was performed at an infection ratio of 1:0.8, corresponding to ratio between the area of the cell layer from which the inoculum originated and the area of the cell layer inoculated. Specifically, 300 µl of medium replaced by 300 µl of WhSBV inoculum. Infections were conducted in three independent replicate wells for each time point. Cell pellets were harvested at 0, 4, 8, 24, 48, 72, 96, 120, 144, 168, 192, 216, and 240 hours post-infection (hpi). At each time point, supernatants were discarded, and cell pellets were collected by trypsinisation using ATV. RNA was extracted using the NucleoMag VET kit as described previously. The extracted nucleic acids were used for RT-qPCR analysis as described above, using specific primer/probe mixes for WhSBV G and ACTB. A DeltaDelta Cq (ΔΔCq) analysis was conducted using R (version 4.3.1; [16]) to compare different inoculation methods, with ACTB serving as the reference gene and Method 1 as the reference method. To ensure consistency, a single batch of inoculum prepared by Method 1 was used throughout most assays.

### Experimental infection of different cell cultures with WhSBV

Bony fish cells RT/F (CCLV no. 0088), CaPi (0112), EPC (0173), RTG-2/f (0686), CCB (0816), SAF-1 (0826), CHSE-214 (1104), GTS-9 (1388), ZF4 (1492), and TiB (1550), the reptilian cells SKL-R (0484), VH-2 (1092), and CDSK (1536), the avian cells QM7 (0466) and DF-1 (1529), and the mammalian cells Vero (0015) and C6 (1452) were obtained from CCLV (see Supplementary Table S3 for a complete list of cell lines). All fish and reptile cell lines were grown in non-vented T25 flasks (Corning) to 95-100% confluence in an appropriate 10 ml growth medium supplemented with 10% FBS. Incubation temperatures were set at 20°C for the fish cells CHSE-214 and RTG-2/f, 26°C for all other fish cells and reptilian cell SKL-R, and 28°C for the reptile cells VH-2 and CDSK. Mammalian and avian cells were grown in vented T25 flasks at 37°C in a humidified atmosphere containing 5% CO_2_.

To initiate the infection experiment, cells were grown to confluence in T25 flasks (25 cm²), as described above. Prior to inoculation (T0), 1 ml of supernatant was collected from each cell culture. The cells were then inoculated with 1 ml of WhSBV inoculum, prepared using the optimized protocol involving three freeze/thaw cycles, as previously described. The inoculum was applied at a of 1:1.7, as described above. Cultures were then incubated for 1 h under the above conditions. After this incubation, a further 1 ml of supernatant was collected (T1). Cells were then incubated and split at regular intervals of 2-3 days at a 2-fold splitting factor for a total of 4-6 passages (P1-6). The splitting process involved collecting the supernatant before each passage, washing and detachment of the cells with ATV, and collecting half of the detached cells for analysis. The remaining cells were used for further culture by adding fresh growth medium supplemented with 10% FBS. The collected supernatants and cell pellets were stored at −80°C for subsequent RNA extraction and RT-qPCR analysis. Negative controls were treated in the same way as infected samples, except that the WhSBV inoculum was replaced by 1 ml RNase-free water (Qiagen). The infection experiment was performed in two independent runs for each cell line. As the infection of reptile, avian and mammalian cell lines was performed separately from the comparison of different fish cell lines, the cypriniform cell line GTS-9 was used as a positive control to validate the infectivity of the inoculum.

Nucleic acid extraction from supernatant and cell pellet samples was performed using the NucleoMag VET kit according to the manufacturer’s instructions. RT-qPCR was then performed using WhSBV G gene and ACTB specific RT-qPCR primer/probe mixes, as described earlier.

### Electron microscopy

WhSBV-infected and uninfected ZF4 cells (see Supplementary Table S3) from the previous infection experiment, and persistently WhSBV-infected GCSB1441 cells were transferred to non-vented T75 flasks for electron microscopy analysis. Cells were fixed in 2.5% glutaraldehyde buffered in 0.1 M sodium cacodylate pH 7.2 (SERVA Electrophoresis). 1% aqueous OsO_4_ was used for post-fixation and 2.5% uranyl acetate in ethanol for *en bloc* staining (SERVA Electrophoresis). After a graded dehydration in ethanol, the samples were cleared in propylene oxide and infiltrated with Glycid Ether 100 (SERVA Electrophoresis). For polymerisation, samples were incubated at 60°C for 3 days. Ultrathin sections were transferred to formvar-coated nickel grids (slot grids; Plano). All grids were counterstained with uranyl acetate and lead citrate before examination on a Talos F200i transmission electron microscope (FEI) at an accelerating voltage of 80 kV.

### Specific detection of viral RNA using RNA *in situ* hybridisation

The cypriniform cell lines (EPC and CCB), the non-cypriniform fish cells (RTG-2/f and TiB), the mammalian Vero line, and the avian DF-1 cells (see Supplementary Table S3) were maintained in non-vented T75 flasks in a suitable medium supplemented with 10% FBS until they reached 95-100% confluence. Cultures were maintained under the conditions described in the previous infection experiment. For infection, 3 ml of medium was replaced with an equal volume of the inoculum prepared via three freeze/thaw cycles, to achieve an infection ratio of 1:1.7, as previously described. For cypriniform and non-cypriniform fish cells were harvested from two independent experiments at early and late time points: 0, 4, 8, 24, 48 hpi for the early time points and 0, 72, 96, 120 and 240 hpi for the late time points. In a separate experiment, the Vero and DF-1 cell lines were harvested exclusively at 0 and 240 hpi, alongside the CCB cell line, which served as a positive cypriniform control. All cells were harvested using a cell scraper. The harvested cells were collected in 50 ml Falcon tubes, centrifuged at 3500 rpm for 15 minutes and the supernatants discarded. The cell pellets were then fixed in 10% neutral buffered formalin (Carl Roth) for at least 24 h. After paraffin embedding, 4-μm thick sections were processed for subsequent RNA *in situ* hybridisation (RNA ISH) using RNAScope 2-5 HD Reagent Kit Red and custom-designed probes targeting WhSBV (-)RNA (genomic RNA; WhSBV-L-O2, catalogue number 1567161-C1, Advanced Cell Diagnostics, Hayward, USA) on all samples according to the manufacturer’s instructions. In addition, custom-designed probes targeting WhSBV (+)RNA (mRNA and antigenomic RNA; WhSBV-G-O1, catalogue number 1567171-C1) were used on CCB samples. As technical controls, probes against the genes for peptidylprolyl isomerase B (PPIB) and dihydrodipicolinate reductase (DapB) were included in each run. In addition, routine staining using hematoxylin-eosin was performed on all samples taken at 0 and 240 hpi respectively for histopathologic evaluation. All slides were scanned using a Hamamatsu S60 scanner and evaluated using NDPview.2 plus software (version 2.8.24, Hamamatsu Photonics, K.K. Japan) by a board-certified pathologist (AB, DiplECVP). The abundance of infected cells was recorded as a percentage in a 10 × 10 grid (size 100 × 100 µm per grid cell). Thus, a positive signal within a grid cell resulted in one percentage point.

In a separate kinetics experiment to correlate with the RNA ISH results, the same fish cell lines as above were seeded in 12-well plates (3.8 cm^2^ cell layer per well) and maintained to 95-100% confluence in appropriate medium supplemented with 10% FBS. Fish cultures were maintained at 26 or 20°C (for RTG-2/f). Inoculation was performed at a 1:0.8 infection ratio as described above, with 300 µl of medium replaced by 300 µl of WhSBV inoculum. Infection of each cell line was performed in three independent replicate wells for each time point. Cell pellets were harvested at 0, 4, 8, 24, 48, 72, 96, 120, 144, 168, 192, 216 and 240 hpi. At each time point, supernatants were discarded and cell pellets were collected by trypsinisation using ATV. RNA was extracted using the NucleoMag VET kit as described previously. RT-qPCR was then performed using G gene and ACTB specific RT-qPCR primer/probe mixes.

### Gene expression analysis of early and late WhSBV infection

The CCB cells (see Supplementary Table S3) were used to study the effect of WhSBV on cellular gene expression. Cells were seeded in 6-well plates (9.6 cm^2^ cell layer per well) (Corning) and cultured for 24 hours to 95-100% confluence in 10% FBS supplemented medium. They were maintained under the conditions described in the previous kinetics experiment. Infection with WhSBV was performed at a 1:1.3 ratio as described above by replacing 500 µl of culture medium in each well with 500 µl of WhSBV inoculum prepared via three freeze/thaw cycles. Negative controls were inoculated with 500 µl of the same medium used for persistently WhSBV-infected GCSB1441 cultivation, replacing 500 µl of the cell culture medium. Cell pellets were collected in two independent experiments to assess early and late infection stages. In the early infection experiment, mock-infected and WhSBV-infected cell pellets were collected at 4, 8, 24 and 48 hpi. For the late-stage experiment, samples were collected at 72, 96, 120 and 240 hpi. In addition, pre-infection cell pellets were collected at 7 and 24 hours before inoculation, the latter representing the 0 hpi time point. Infection was performed in three independent replicate wells for each time point. Cell pellets were collected by trypsinisation after supernatants were discarded. Total RNA was extracted using Trizol, chloroform and the Agencourt RNAdvance Tissue Kit as previously described. RT-qPCR was performed on all samples using G-gene and ACTB-specific RT-qPCR primer/probe mixes to validate samples prior to sequencing. Transcriptomic libraries were prepared as previously described, but including ERCC spike-in mix (Invitrogen) as an internal control. Subsequently, the libraries were pooled and sent for sequencing on an Illumina NovaSeq 6000 system (Novogene GmbH) running in paired-end 150 bp mode. Raw reads were subsequently trimmed using *TrimGalore!* (v0.6.10; [13]) and then analysed using the nf-core “RNAseq” pipeline (version 3.14.0; [18, 19]). In detail, the common carp reference genome GCF_018340385.1 was used for splice-aware mapping with STAR and subsequent quantification with salmon. The quantification data was then further analysed in R (version 4.3.1; [16]) and differentially expressed genes (DEGs) were deduced using the DESeq2 package (v1.40; [20]). Log2-fold changes were shrinked using the function “lfcShrink” with the “apeglm” shrinkage estimator [21]. The cut-off values for DEGs were set to absolute shrinked log2-fold change > 1 and adjusted *p-value* < 0.05. To explore global gene expression patterns, principal component analysis (PCA) was performed using R (version 4.3.1; [16]). Transcriptomic data have been deposited in ArrayExpress under E-MTAB-14894.

### Prediction of regulatory and splice sites in WhSBV genome

The trimmed raw reads from the metagenomic RNA datasets from persistently WhSBV-infected GCSB1441 and GCSB1542 cells (see Supplementary Table S1) were mapped back to the RACE corrected WhSBV genome (PV171101) using the “bbmap” function from BBTools (version 39.33; [22]). Duplicates were removed from mapped reads using the “MarkDuplicates” function from Picard toolkit (version 2.20.4; [23]) and reads were separated by mapping orientation using Samtools (version 1.22.1; [24]). The genome coverage for forward and reverse deduplicated reads was obtained using Bedtools (version 2.31.1; [25]) und was plotted using R (version 4.3.1; [16]). Poly(A)-addition sites were deduced using the same dataset and reference with ContextMap (version 2.7.9; [26]) and bowtie (version 1.3.1; [27]). Potential introns were detected using STAR (version 2.7.11b; [28]) running in “2-pass” mode, enabling sensitive novel junction discovery. The function “FIMO” from the MEME Suite (v5.5.2; [29]) was used to identify the conserved motif “`AKUUAAYAAAAACAUGAA`” [2] of potential transcription termination and start sites within the WhSBV genome sequence, respectively.

### Validation of predicted splice sites in WhSBV genome

To experimentally validate each predicted splice site, primers were designed to amplify the specific genomic regions by RT-PCR. Purified RNA was extracted by RNeasy Mini kit as described above. cDNA from poly(A) RNA was synthesized using the Protoscript II First Strand cDNA Synthesis Kit and oligo-dT primers (New England Biolabs). The cDNA products were then digested with an RNase cocktail and purified using AMPure XP beads. Conventional Taq-PCR was performed using the Accuprime *Taq* DNA Polymerase System (Invitrogen) with primers specific for the predicted splice sites within the viral M and L genes (see Supplementary Table S2). RNase-free water was used instead of cDNA as a negative control. The amplified DNA products were run on a 2% agarose gel and bands were excised. The DNA was extracted using a QIAquick Gel Extraction Kit (Qiagen) and then sequenced using the BigDye Terminator v1.1 Cycle Sequencing Kit (Applied Biosystems) on a 3500 Genetic Analyzer (Applied Biosystems).

### Specific detection of viral RNA using Northern blot

RNA was extracted from the persistently WhSBV-infected GCSB1441, WhSBV-infected and uninfected EPC cells (see Supplementary Tables S1 and S3) from the previous infection experiment using the TRIzol-chloroform method and the RNeasy Mini Kit with DNase I (Qiagen), as previously described. RNA quality and quantity were assessed using the TapeStation System 4150 (Agilent Technologies) and NanoDrop Lite spectrophotometer as described previously. Subsequently, 2-4 µg of total RNA was glyoxylated using Glyoxal (Sigma-Aldrich) at 56°C for 45 min before separation on a 0.9% denaturing agarose gel (4.7% formaldehyde, Carl Roth) in a phosphate buffer system. RNA was transferred overnight to Hybond-N membranes (Cytiva Amersham) using SSC transfer buffer. RNA was cross-linked to the membrane and viral RNA (mRNA and genomic RNA) was detected using DNA probes. Probes were generated by RT-PCR using specific forward and reverse primers against each gene of WhSBV (see Supplementary Table S2) using the Platinum Script III One-step RT-PCR Kit (Thermo Fisher Scientific) according to the manufacturer’s instructions. The amplified cDNA products were then run on a 2% agarose gel. After amplification, the products were purified using AMPure XP beads according to the manufacturer’s protocol. The quantity of purified products was measured using a NanoDrop Lite spectrophotometer and labelled with ^32^P-CTP (Hartmann-Analytic) using a Nick translation kit (GE Healthcare). Hybridisation was performed overnight at 60-65°C and the signal was analysed after washing using the CR35 phosphoimager system (Dürr Medical).

### Specific detection of WhSBV proteins

Approximately 3×10^5^ persistently WhSBV-infected GCSB1441 cells were harvested, resuspended in 1× LDS Buffer (Thermo Fisher Scientific) containing 100 mM DTT (Sigma-Aldrich) and heated for 10 min at 70°C shaking at 1400 rpm in a Thermomixer (Eppendorf). Reconstitution buffer (50 mM HEPES/KOH, pH 7.5, 100 mM NaCl, 1 mM EDTA, pH 8.0, 0.5% [w/v] Sodium deoxycholate, 1% [v/v] Triton X-100) was added to a final volume of 100 µl. Samples were reduced in 10 mM DTT for 30 min at 60°C followed by alkylation in 25 mM iodoacetamide (Sigma-Aldrich) for 25 min at RT in the dark with orbital shaking at 800 rpm in a Thermomixer (Eppendorf). The reaction was quenched by incubation with 10 mM DTT f.c. for 5 min at RT. For protein binding 125 µg hydrophilic and 125 µg hydrophobic Sera-Mag Carboxylate-Modified Magnetic Particles (GE Healthcare) together with 150 µl absolute ethanol (Carl Roth) were added to each sample and incubated for 15 min at RT at 1000 rpm in a Thermomixer (Eppendorf). Beads were washed three times with 80% LC/MS grade ethanol (Supelco). Residual ethanol was removed prior to adding 1 µg LC-grade trypsin (Serva) in 100 µl of 50 mM ammonium bicarbonate buffer pH 8. After incubation for 4 h at 37°C and 1000 rpm in a Thermomixer (Eppendorf) peptides were collected and desalted on StageTips [30]. The amount of 400 ng digested peptides were analysed with a nanoElute2 HPLC system (Bruker) coupled to a TimsTOF HT mass spectrometer (Bruker). Peptides were separated on an Ultimate CSI 25×75 C18 UHPLC column (Ionopticks) by running an optimized 100 min gradient of 2 to 95% MS-grade acetonitrile with 0.1% formic acid at 250 nl/min at 50°C. The mass spectrometer was operated with the manufacturer provided DDA_PASEF_1.1sec_cycletime method. Mass spectrometry raw files were processed with MaxQuant (version 2.4.13.0) [31] using a custom virus database (6 entries) and the *C. idella* database (C_idella_female_genemodels.v1; 32,811 entries) with standard settings.

## RESULTS

### Metagenomic screening of different fish cell lines reveals presence of WhSBV

During metagenomic sequencing of 48 fish cell lines, the cypriniform cell lines GCSB1441 and GCSB1542 were found to contain sequences matching WhSBV (Supplementary Figure S1). Complete viral genomes were assembled from both cell lines, sharing 99.9% nt identity with each other and 99.9% and 87.8% nt identity with WhSBV variants CIK (BK063520) and DSYS4497 (NC_055169), respectively. Both currently published WhSBV reference genomes (NC_055169 and BK063520) were identified through metagenomic sequencing and *de novo* assembly. The NC_055169 genome was 8,989 nt in length, while the BK063520 variant was 8,985 nt. However, 3’ and 5’ RACE extended the WhSBV genome to 9,008 nt (Supplementary Figure S2). These RACE results did not alter the internal genome sequence but revealed additional nucleotides beyond those captured in the metagenomic assembly. Compared to the current reference genome sequence in Genbank (RefSeq NC_055169), the complete WhSBV genome sequence presented here is 7 and 11 nt longer at the 3’ and 5’ ends, respectively. This observation highlights that metagenomic sequencing alone may not capture the full-length viral genome, and complementary methods such as RACE are essential to accurately determine the complete genome ends.

Based on the complete WhSBV genome sequences, gene-specific RT-qPCR assays targeting the N, P, G, M, and L gene regions were established. All five assays were evaluated using total RNA from the persistently WhSBV-infected GCSB1441 and GCSB1542, and uninfected GK cells as control. The Cq value of the individual WhSBV gene specific RT-qPCRs were comparable in GCSB1441 and GCSB1542, while no amplification was observed in GK cells (see Supplementary Figure S3). Among the targets, the G gene-specific RT-qPCR assay demonstrated the highest sensitivity and consistently produced the lowest Cq values. It was therefore selected for further screening and for subsequent analyses. Using serial dilutions, a Cq cut-off value of 37 was defined for the G gene-specific RT-qPCR, corresponding to the highest dilution that consistently produced detectable amplification.

Subsequent testing of the 48 cell lines used for metagenomic analysis with the G gene assay identified WhSBV RNA in GCSB1441 and GCSB1542 cell lines, respectively, while all other cell lines tested negative (see Supplementary Table S1). Furthermore, qPCR assays targeting the WhSBV N, P, G, M and L gene regions without prior reverse transcription were negative for GCSB1441 and GCSB1542 cells, indicating that the detected WhSBV sequences likely represent genuine viral RNA rather than endogenous viral elements (EVE) integrated into the host DNA genome (see Supplementary Figure S3).

### Cell lysis is necessary for efficient release of WhSBV viral particles

Five inoculum preparation methods were compared to assess their ability to induce WhSBV replication in CCB cells (Supplementary Table S3). In the inoculated cultures, WhSBV and cellular ACTB RNA levels were monitored using RT-qPCR over a time period of 240 hpi (**Figure 1A**). Three cycles of freeze/thawing without subsequent centrifugation appeared to yield the highest amounts of infectious virus, as higher viral RNA loads were detectable in the inoculated cells already at early time points, as compared to methods employing sonification with or without freeze/thaw cycles or hypertonic treatment (**Figure 1A**). For these latter methods, cell debris was removed by centrifugation. Cells infected with untreated supernatant barely showed any increase of viral RNA over time, suggesting that the measured viral RNA might have been residual inoculum rather than result of an active viral replication (**Figure 1A**). Based on its performance, consistency, and practical ease, freeze/thaw lysis was selected for use in subsequent infection experiments.

**Figure 1:**
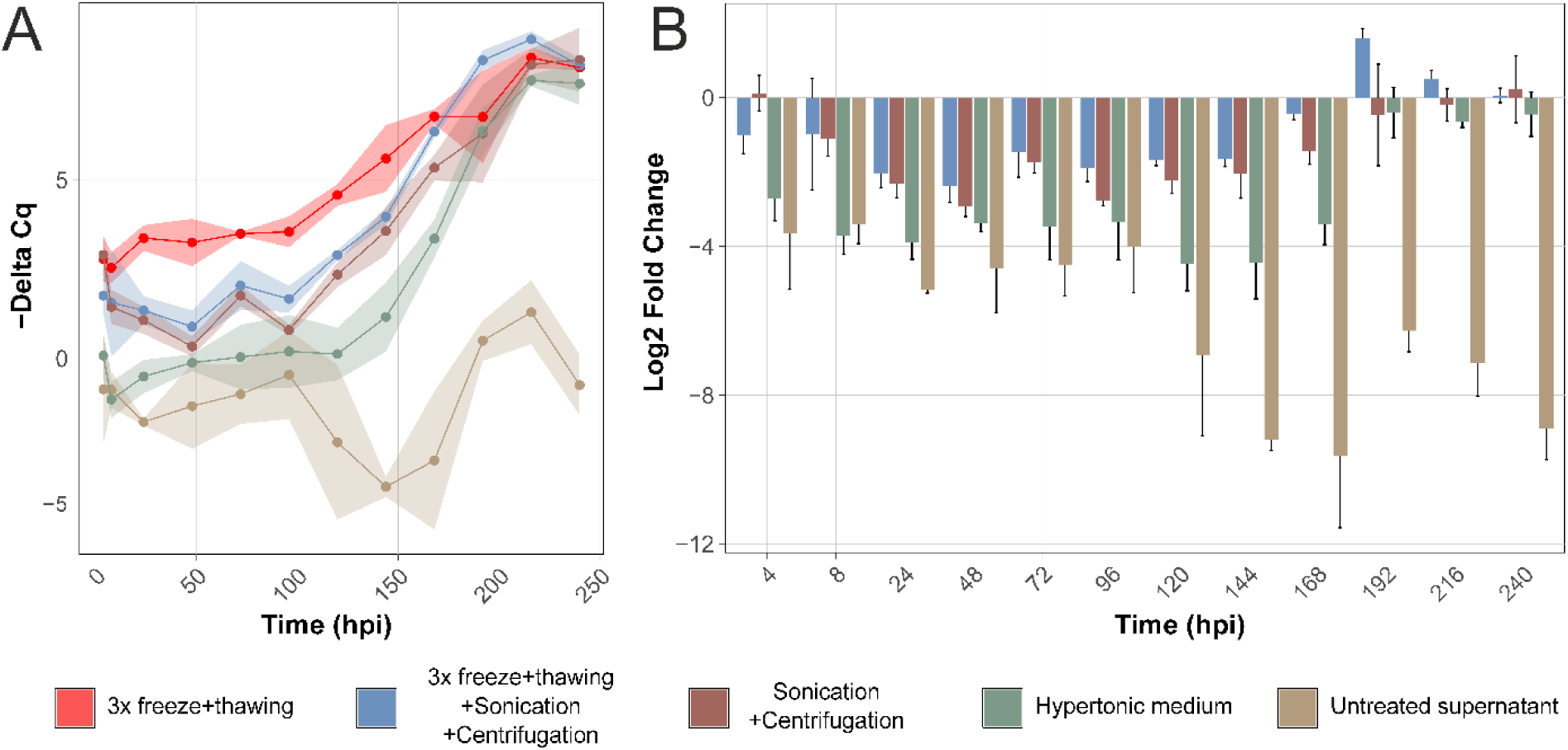
Evaluation of inoculum preparation for WhSBV propagation in CCB cells. (**A**) CCB cells were experimentally infected with WhSBV preparations, which had been produced from persistently WhSBV-infected GCSB1441 cells using five different methods (each represented by colour). Cell pellets were harvested at 0, 4, 8, 24, 48, 72, 96, 120, 144, 168, 192, 216, and 240 hours post infection (hpi) and RNA levels of the viral glycoprotein (G) and cellular beta actin (ACTB) genes were detected by reverse transcription quantitative PCR (RT-qPCR). Values are presented as negative “Delta Cq”. (**B**) Relative change of viral RNA levels in CCB cells over time in comparison to “3x freeze/thawing” method. Values are presented as log_2_ fold changes calculated using the “Delta Delta Cq” method. Each point/bar represents the mean of three technical replicates (± standard deviation).

### WhSBV establishes persistency only in cypriniform fish cells

In order to test the infectivity of WhSBV and its potential *in vitro* host range, we used cell lysate from persistently WhSBV-infected GCSB1441 cells to inoculate various cell lines from bony fish, reptiles, birds and mammals (see Supplementary Table S3). Following inoculation, the cells were propagated for several passages, and both cell pellets and supernatants were tested for presence of viral RNA using specific RT-qPCR.

WhSBV RNA was detected in the supernatant and cell pellet of all fish cell lines tested during the first one to two passages. In cells from the order Cypriniformes, including cyprinid (CaPi, CCB and GTS-9), leuciscid (EPC) and danionid (ZF4) species, viral RNA was retained over all six consecutive passages. The amount of viral RNA detected started to increase compared to freshly inoculated cells (T1) and eventually reached a plateau at around Cq 22 in both supernatant and cell pellet. This stabilisation occurred between passages 2 and 4 for the CaPi, EPC, CCB and GTS-9 cell lines. For zebrafish cell line ZF4, the amount of viral RNA initially decreased after inoculation and passaging until passage 2-3, but then recovered, reaching Cq values of about 27 in supernatant and cell pellets (**Figure 2A**).

**Figure 2:**
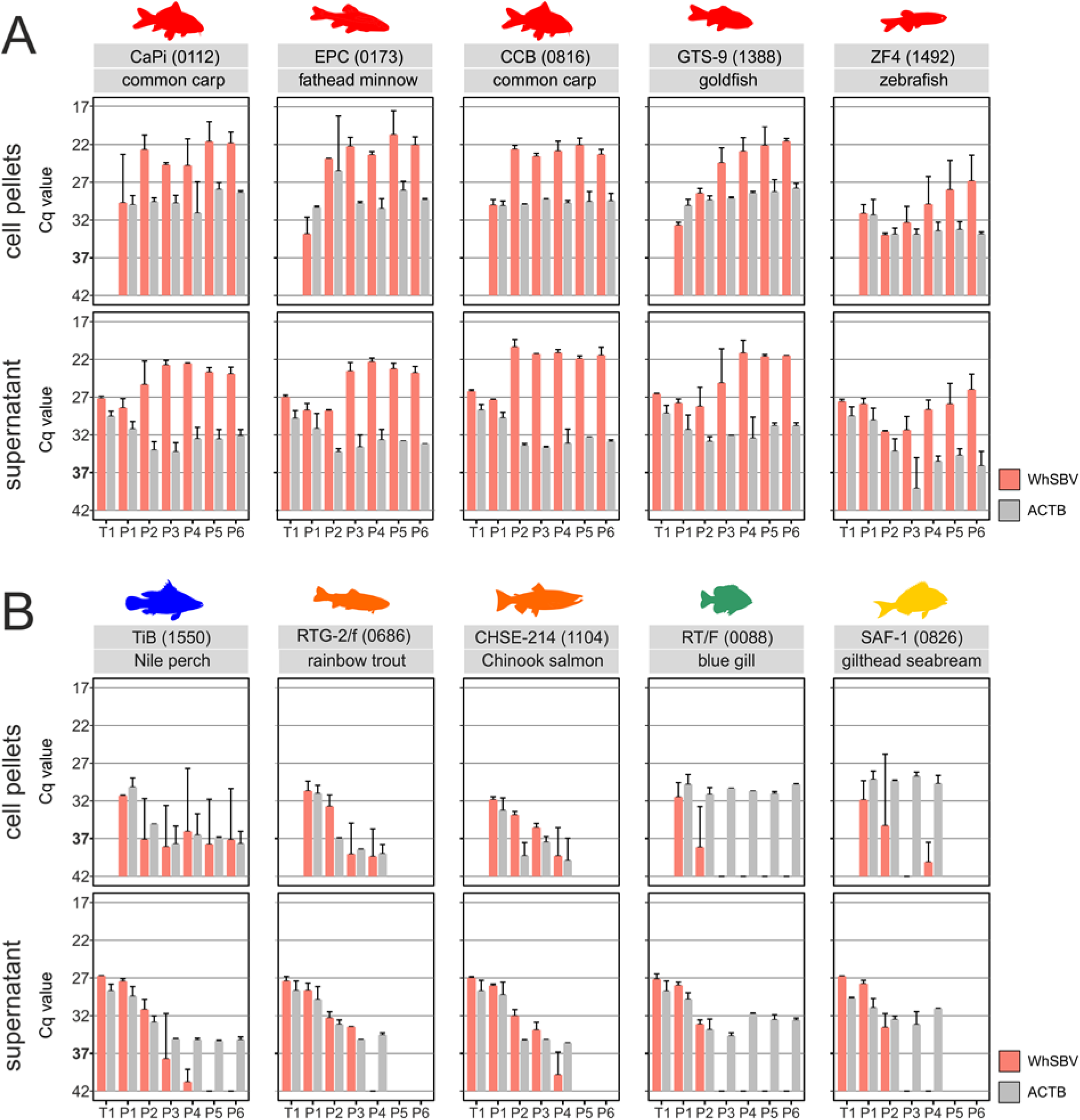
WhSBV RNA levels over successive passages in different fish cell lines. Different fish cell lines were inoculated with WhSBV. The cells were then passaged for 4-6 times by splitting. (**A**) Cell lines from the taxonomic order of Cypriniformes (red); (**B**) Cell lines from the taxonomic orders of Cichliformes (blue), Salmoniformes (orange), Centrarchiformes (green) and Spariformes (yellow). RNA levels of viral glycoprotein (WhSBV; red bars) and cellular beta actin (ACTB; grey bars) genes were detected by specific RT-qPCR in cell pellets (top row) and cell culture supernatants (bottom row) from each passage. Results are presented as the arithmetic mean (± standard deviation) of the cycle of quantification (Cq) values of two independent experiments. A Cq cut-off value of 37 was set as limit of detection for WhSBV.

In contrast, the non-cypriniform fish, centrarchid (RT/F), and sparid (SAF-1) showed no evidence of WhSBV replication after inoculation (**Figure 2B**). Immediately after inoculation (T1), viral RNA levels in the supernatant were similar to those of the cypriniform cell lines (**Figure 2A**), but dropped below the detection limit in cell pellets and supernatant after passage 2 (**Figure 2B**). Although, viral RNA was detected in SAF-1 cell pellets at passage 4, the levels remained below the defined Cq cut-off and were not indicative of active replication. The cichlid (TiB) and salmonid (RTG-2/f and CHSE-214) cell lines exhibited reduced cell growth over the course of experiment, which was reflected by a decline in the measured ACTB RNA levels, a trend also observed in corresponding uninfected controls. Viral RNA remained detectable in cell pellets until the end of the experiment up to passage 6 for (TiB) and passage 4 for (RTG-2/f and CHSE-214), but without a detectable increase in levels. In several instances, Cq values fluctuated near or fell below the established detection cut-off, indicating the absence of sustained viral replication (**Figure 2B**).

In the reptilian cell lines SKL-R, VH-2, and CDSK, WhSBV RNA levels in the culture supernatant decreased after inoculation but remained detectable up to passage 6, with Cq values stabilizing around 28 and 35, however, viral RNA levels fell below the defined Cq cut-off at passage 5 and 6 in SKL-R (**Figure 3**). In the SKL-R cell pellets, viral RNA was still detectable up to passage 6, although consistently at low levels, yet above the cut-off. In contrast, CDSK and VH-2 pellets showed an increase in viral RNA levels during passages 5 and 6, reaching Cq values of 28 and 30, respectively. Decline in ACTB RNA levels was also observed in the reptilian cell lines, mirroring the trend in cichlid and salmonid cell lines. Similar decreases in ACTB expression were also noted in the corresponding uninfected controls.

**Figure 3:**
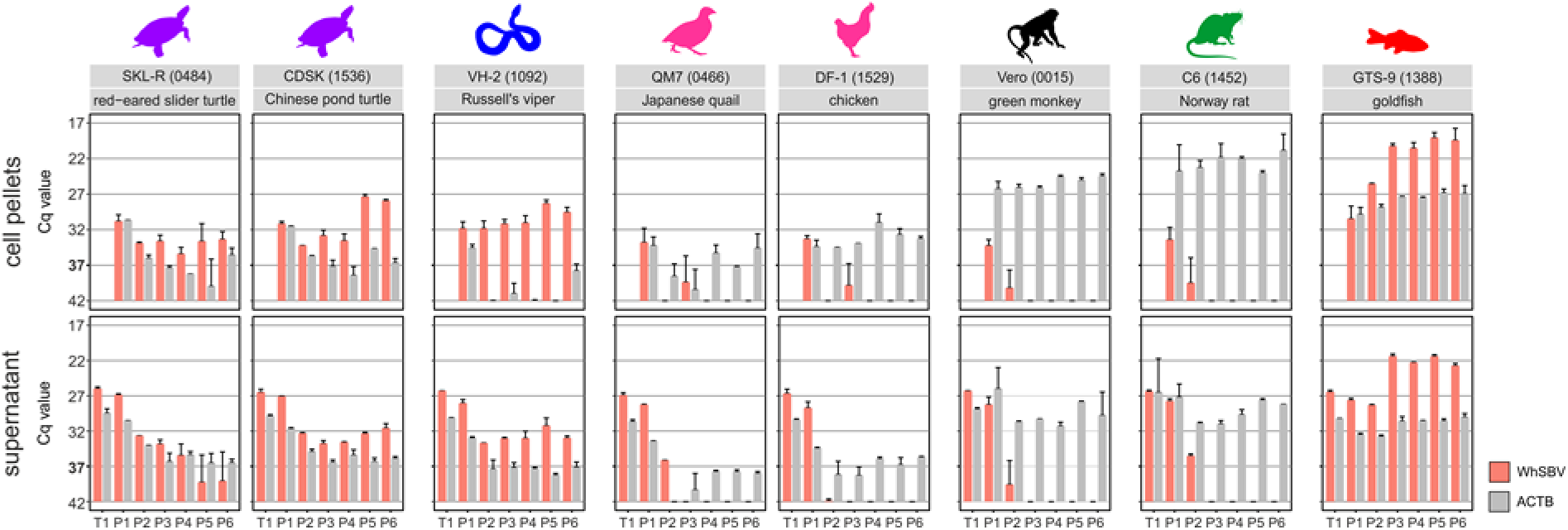
WhSBV RNA levels over successive passages in different non-fish cell lines. Different cell lines from the taxonomic orders of Testudines (purple), Squamata (blue), Galliformes (pink), Primates (black), and Rodentia (green) were inoculated with WhSBV and subsequently passaged for 6 times by splitting of the cells. The cypriniform fish cell line GTS-9 (red) was used as a positive control to validate the infection experiment. RNA levels of viral glycoprotein (WhSBV; red bars) and cellular beta actin (ACTB; grey bars) genes were detected by specific RT-qPCR in cell pellets (top row) and cell culture supernatants (bottom row) from each passage. Results are presented as the arithmetic mean (± standard deviation) of the cycle of quantification (Cq) values of two independent experiments. A Cq cut-off value of 37 was set as limit of detection for WhSBV.

In the mammalian Vero and C6 cell lines and the avian QM7 and DF-1 cell lines, WhSBV RNA levels in the cell pellets or supernatants dropped below or near the detection limit already after passage 1-3, respectively (**Figure 3**).

All negative controls used in this experiment showed no Cq values for the viral glycoprotein.

No apparent cytopathic effect was observed in any of the cell cultures used throughout the experiment.

Using transmission electron microscopy, putative virus particle-like structures were observed in WhSBV-inoculated ZF4 cell lines, whereas similar structures were absent in non-inoculated controls. These structures appeared spherical to pleomorphic without visible peplomers and a diameter of approximately 150-180 nm (**Figure 4A**). Potential membrane associated budding was rarely observed at the plasma membrane of the persistently WhSBV-infected GCSB1441 cells (**Figure 4B**).

**Figure 4:**
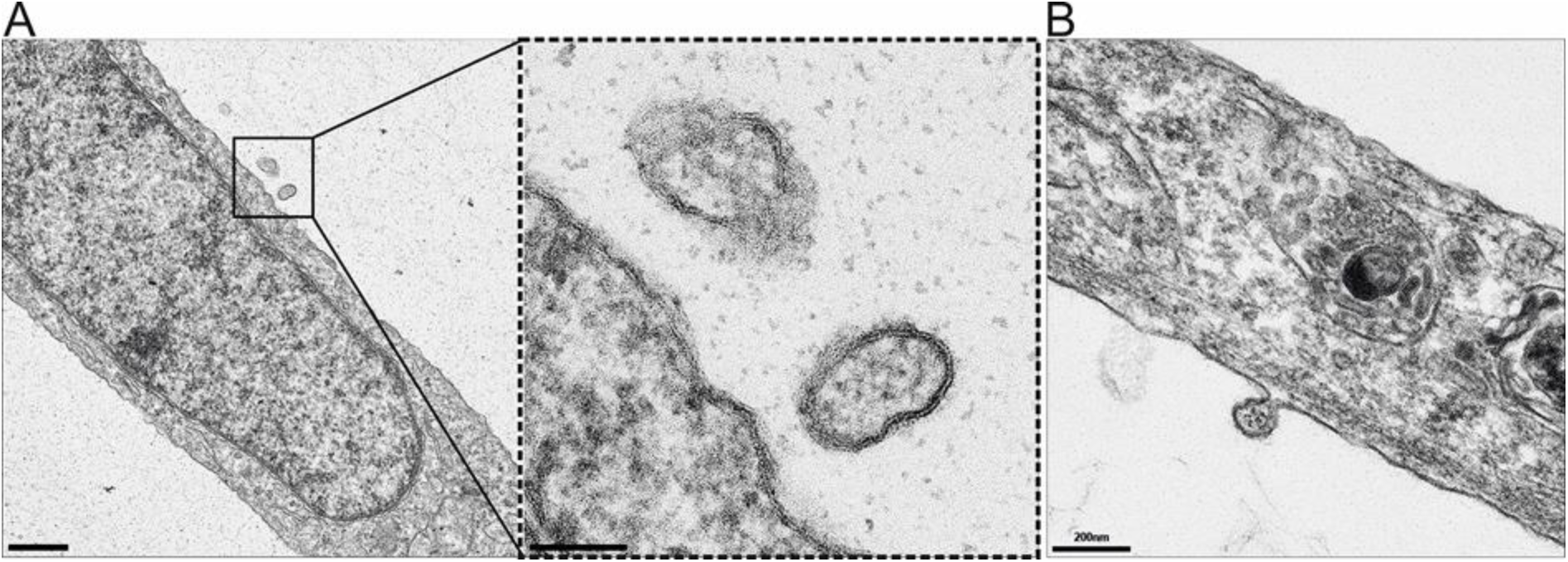
Detection of putative WhSBV-like particles by transmission electron microscopy. (**A**) Putative virion-like structures observed in the experimentally infected ZF4 cell lines. These structures appeared spherical to pleomorphic, with diameters ranging from approximately 150-180 nm. (**B**) Rare membrane-associated budding events were observed at the plasma membrane of persistently WhSBV-infected GCSB1441 cells, which may represent viral budding.

### WhSBV replicates efficiently only in cypriniform cell lines

In order to gain deeper insights into viral propagation in different fish cell lines, a kinetic infection experiment was performed over 10 days using both cypriniform (EPC and CCB) and non-cypriniform fish cells (RTG-2/f and TiB). The level of viral RNA load was assessed by RT-qPCR and the presence of viral RNA at the cellular level was assessed by RNA ISH at different time points.

In the cypriniform EPC and CCB cell lines, viral RNA levels increased exponentially until 12 hpi, reaching Cq values of about 18.6 and 19.5 for EPC and CCB, respectively. In contrast, the viral RNA levels in salmonid RTG-2/f and cichlid TiB cell lines showed stagnation or linear increase (**Figure 5A**). The presence of viral (-)RNA was detected by specific RNA ISH in both cypriniform (EPC and CCB) and non-cypriniform fish cell lines (RTG-2/f and TiB) from 4 hpi onward. In detail, RNA ISH confirmed the RT-qPCR results from the kinetic experiment, as the number of infected cells, indicated by the presence of viral (-)RNA, increased in a time-dependent manner for the cypriniform cell lines EPC and CCB. At early time points (4 - 72 hpi), infected single cells appeared, followed by foci of positive cells, later confluent and finally diffuse labelling of the cell pellet (**Figure 5B**). Percentage quantification of individual cell pellets for EPC showed widespread infection, with 95% of the fields analysed showing positive detection as early as 4 hpi. Infection remained widespread (63-100% of infected test fields) throughout the course of the experiment until 240 hpi (see Supplementary Figure S4). In CCB cells, only 29% of the test fields were infected at 4 hpi and longer cultivation correlated with a higher proportion of positive test fields, eventually reaching 100% at 96, 120 and 240 hpi (see Supplementary Figure S4). Similar observations were made using a custom-designed probe against WhSBV (+)RNA on CCB cell lines (Supplementary Figure S5). In terms of labelling signal, infected cells harvested at early time points were more likely to show fine-granular cytoplasmic labelling, and larger, globular signals were consistently found at 48 hpi and later (**Figure 5B**). In clear contrast, the infection pattern in the non-cypriniform fish cell lines were essentially confined to single cells and showed no evidence of increasing infection abundance over time (**Figure 5B**). Only the intensity of the chromogen signal within the cell increased in TiB from 72 hpi, but in RTG-2/f this phenomenon was limited to 72 and 96 hpi. RTG-2/f and TiB cells ultimately yielded 28% and 16% infected test fields at 240 hpi, respectively (see Supplementary Figure S4).

**Figure 5:**
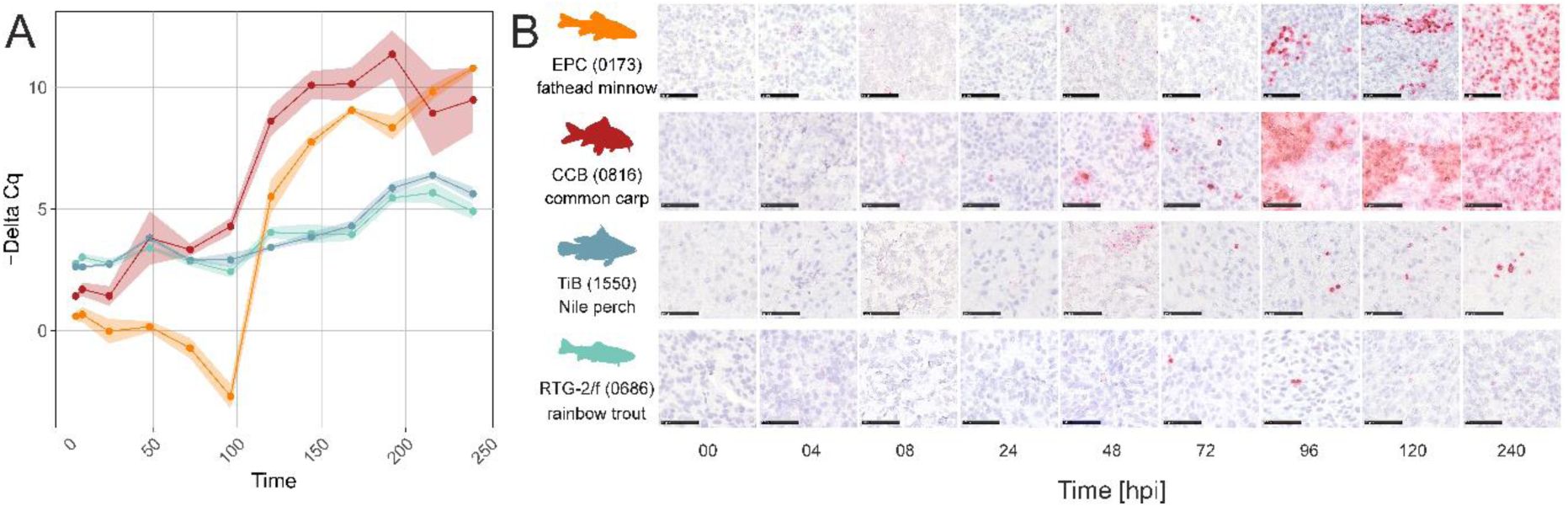
**Detection of viral RNA by RT-qPCR and RNA *in situ* hybridisation**. (**A**) RNA levels of the viral glycoprotein (WhSBV) and cellular beta actin (ACTB) genes were detected by reverse transcription quantitative PCR (RT-qPCR) and are presented as negative Delta cycle of quantification (-Delta Cq) values for RNA extracted from cell pellets of EPC (orange), CCB (red), RTG-2/f (greenish-blue) and TiB (blue-grey) at different hours post inoculation (hpi). Results are presented as arithmetic means (± standard deviation) of three replicates. A Cq cut-off value of 37 was set as limit of detection for WhSBV. (**B**) Detection of viral RNA by RNA *in situ* hybridisation in formalin-fixed cell pellets from WhSBV-inoculated cypriniform (CCB, EPC) and non-cypriniform (TiB, RTG-2/f) cell lines. One representative image is shown for each cell line and time point. The black scale bar represents 50 µm.

For mammalian Vero and avian DF-1 cell lines, only a single time point at 240 hpi was collected, alongside CCB which used as a positive control. RNA ISH analysis of mammalian and avian DF-1 cell lines showed no detectable WhSBV RNA at either the control 0 hpi or at 240 hpi (data not shown).

Using routine staining (hematoxylin-eosin), neither viral inclusion bodies nor other morphological evidence of a cytopathogenic effect could be identified in EPC, CCB, RTG-2/f, TiB, Vero, and DF-1 cell lines (see Supplementary Figure S6).

### Cellular response to early and late WhSBV infection is limited in cypriniform cells

To investigate cellular transcriptional changes associated with WhSBV infection, we performed RNA sequencing (RNAseq) of RNA transcripts at early and late time points following viral inoculation in CCB cells (**Figure 6A**). Quantitative RT-qPCR and sequencing confirmed the presence of WhSBV RNA and the onset of infection at 4 hpi, with viral RNA levels progressively increasing at all subsequent time points (**Figure 6B**). This trend was consistent with the RNA ISH data, which showed a gradual increase in the percentage of infected CCB cells over time, reaching 100% infection at 96, 120, and 240 hpi (**Figure 6B**).

**Figure 6:**
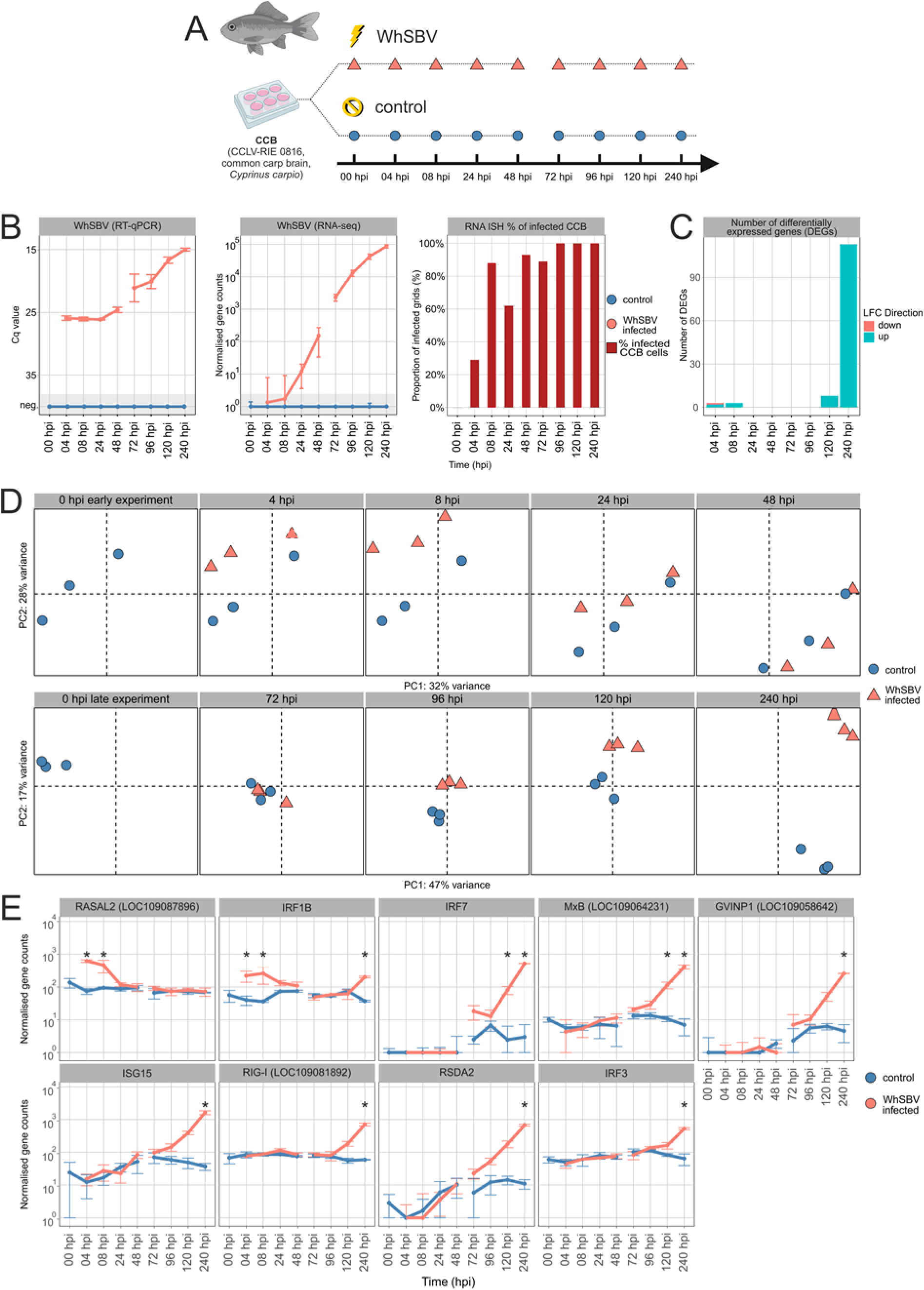
Cellular response to early and late WhSBV infection in the cypriniform brain cells CCB. (**A**) CCB cells were experimentally WhSBV-infected (orange triangles) and mock-infected (blue dots) in two consecutive experiments, covering early (0, 4, 8, 24, and 48 hours post infection (hpi)) and late (0, 72, 96, 120, and 240 hpi) infection stages, respectively. (**B**) Viral RNA was detected by reverse transcription quantitative PCR (RT-qPCR; left panel) and RNA sequencing (RNAseq; middle panel). The RT-qPCR targeted the viral glycoprotein and is presented as cycle of quantification values (Cq), while RNAseq quantified sequence reads corresponding to the WhSBV genome and is presented as normalised gene counts. Arithmetic means (± standard deviation, SD) of three replicate wells are shown for Cq values and normalised gene counts. WhSBV-specific RNA *in situ* hybridisation (RNA ISH) data from a separate experiment are shown for comparison (right panel; see Supplementary Figure S4). (**C**) Differentially expressed genes (DEGs) with an absolute log2-fold change (LFC) of > 1 and an adjusted *p-value* < 0.05 were derived from the RNAseq data using the DESeq2 package (v1.40), and their number and LFC direction are shown for each time point. (**D**) Principal component analysis (PCA) was performed on the RNAseq gene expression data using R (v4.3.1). Orange triangles and blue dots represent replicates of WhSBV-infected and mock-infected CCB cells, respectively. (**E**) Trends in transcript levels of selected genes known to be key antiviral genes in the early innate immune response to viral infection in fish. These genes include RASAL2 (RAS GTPase-activating protein 2-like [LOC109087896]), IRF1B (interferon regulatory factor 1), IRF7 (interferon regulatory factor 7), IRF3 (interferon regulatory factor 3), MxB (interferon-induced GTP-binding protein MxB-like), GVINP1 (interferon-induced very large GTPase 1-like [LOC109058642]), ISG15 (interferon-stimulated gene 15), RIG-I (antiviral innate immune response receptor RIG-I [LOC109081892]), and RSDA2 (viperin, radical S-adenosyl methionine domain containing 2). Asterisks indicate time points with significantly differentially gene expression as compared to uninfected controls. The complete list of DEGs can be found in Supplementary Table S4. Results are presented as the arithmetic/geometric mean (± standard deviation) of three replicate wells.

The number of DEGs remained minimal at early and late time points but became more pronounced at 240 hpi, indicating a delayed transcriptional response to infection (**Figure 6C**). PCA of early time points revealed clear transcriptional differences between WhSBV-infected and control cells at 4 and 8 hpi, which diminished at 24 and 48 hpi, with both groups becoming more similar (**Figure 6C**). In contrast, the PCA of late time points experiment showed no significant transcriptional changes at 72 and 96 hpi, However, minimal transcriptional alterations were observed at 120 hpi, followed by distinct clustering of infected and control samples at 240 hpi, suggesting a change in the host transcriptional response at later stages of infection (**Figure 6D**).

Differential gene expression analysis of early time points identified three DEGs at 4 or 8 hpi. Specifically, RASAL2 (RAS GTPase-activating protein 2-like [LOC109087896]) and IRF1B (interferon regulatory factor 1) were upregulated at 4 hpi and 8 hpi (**Figure 6E**), whereas EDEM3 (ER degradation enhancing alpha-mannosidase-like protein 3 [LOC109095466]) was upregulated exclusively at 8 hpi. Conversely, ABCC1 (multidrug resistance-associated protein 1-like [LOC109090229]) was downregulated at 4 hpi. The expression of these genes returned to baseline levels by 24 and 48 hpi. At later stages of infection (72 and 96 hpi), no DEGs were detected between infected and control cells. At 120 hpi, eight DEGs were upregulated, including IRF7 (interferon regulatory factor 7) and MxB (interferon-induced GTP-binding protein MxB-like) (**Figure 6E**). At 240 hpi, 113 DEGs were upregulated, including IRF7, MxB, and several other key antiviral genes involved in early innate immune responses in fish, such as IRF3 (interferon regulatory factor 3), ISG15 (interferon-stimulated gene 15), and RSDA2 (viperin, radical S-adenosyl methionine domain containing 2), along with other genes involved in interferon pathways, RIG-I (antiviral innate immune response receptor RIG-I [LOC109081892]) and GVINP1 (interferon-induced very large GTPase 1-like [LOC109058642]) (**Figure 6E** and Supplementary Table S4 for complete list of DEGs).

### Detection of multicistronic and spliced WhSBV gene transcripts

To investigate the transcriptional profile and splice sites of the WhSBV RNA, we mapped raw mRNA sequence data from the persistently WhSBV-infected GCSB1441 and GCSB1542 cells to the *de novo* assembled and RACE corrected WhSBV genome (**Figure 7A**). The observed sequence coverage across the genomes was uneven, with abrupt increases and decreases observed in certain potential intergenic regions. These coverage variations corresponded in part to transcription start and termination sites identified by motif prediction (see Supplementary Table S5). In detail, three transcriptional start sites (S1-3) were located upstream of the N, X and M ORFs, respectively. Start sites S1 and S3 were marked by an increase in read coverage, indicating transcription initiation. Four transcription termination sites were detected by motif prediction: T1, T2 and T4 were located downstream of the N, G and L ORFs, respectively, while T3 was found within the L ORF. The termination sites T2 and T3 were characterised by a marked decrease in sequence coverage and termination sites T1-3 coincided with reads transitioning to poly(A)-tails (**Figure 7A**). S2 and S3 were located immediately adjacent to T1 and T2, respectively. As the 5’ RACE analysis was performed on total RNA including mRNA, Sanger sequencing of some 5’ RACE products identified the likely transcription start site S1 at nucleotide position 33 to 35 in the viral genome, marked by the sequence “GAA”, matching the predicted sequence of S2 and S3. Regions of the genome with comparable levels of sequence coverage between adjacent start and termination sites were interpreted as belonging to the same viral RNA transcript or mRNA. We observed no coverage drop at termination signal T1 and the majority of transcripts seem to skip T1, indicating that N, X/P and G genes can be expressed from polycistronic mRNA (**Figure 7A**). However, we observed reads transitioning to poly(A)-tails at T1 that belong to the monocistronic N mRNA. The mRNAs containing the M and L genes shared S3 as common start site. However, their transcript levels differed with the M gene being comparably highly expressed than the L gene (**Figure 7A**).

**Figure 7:**
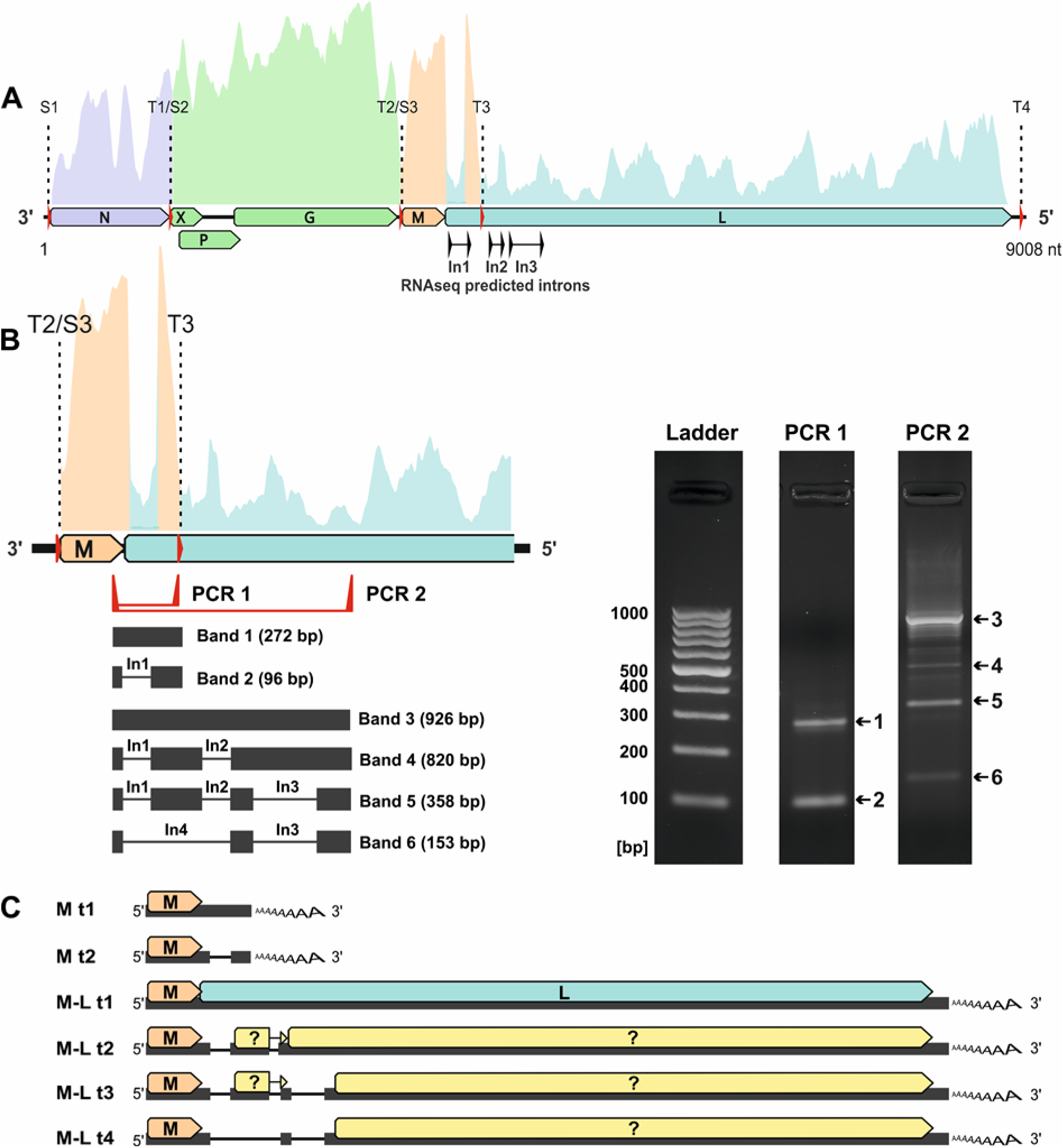
Detection of spliced WhSBV RNA in persistently WhSBV-infected GCSB1542 cells. (**A**) The trimmed raw reads from the metagenomic RNA datasets from persistently WhSBV-infected GCSB1441 and GCSB1542 cells were mapped to WhSBV genome (PV171101) using “bbmap” (version 39.33). In the schematic WhSBV genome, open reading frames (ORFs) are shown as arrows and transcription start sites (S1-S3) and termination sites (T1-4) are indicated by red arrows and dashed lines. ORFs that are likely to be located on the same RNA transcript are shown in the same colour. Potential introns (In1-3) were detected using STAR (version 2.7.11b) and are indicated by black arrows. (**B**) RNA was extracted from the persistently WhSBV-infected GCSB1542 cell line and cDNA was synthesised using oligo-dT primers. A PCR was performed using this cDNA and primers targeting the predicted M and L splice sites (PCR 1 and 2). The PCR products were run on a 2% agarose gel (right panel). Bands 1-6 were excised, purified and sequenced by Sanger sequencing. The sequences were mapped back to the respective WhSBV genome in order to confirm the predicted introns In1-4 (black lines). (**C**) Alternative transcripts of WhSBV were inferred from the detected splice sites, and transcription start and termination sites. The schematic shows six alternative transcripts, designated as M transcripts 1-2 (M t1-t2) or M-L transcripts 1-4 (t1-t4). Arrows indicate possible ORFs.

In addition, mapping of sequence reads revealed the presence of three potential introns (In1-3) located at the beginning of the L ORF. Conventional RT-PCR and Sanger sequencing confirmed the presence of In1-3 within the L ORF and also revealed an additional, unpredicted splice site, In4, within the L ORF (**Figure 7B**). Based on the above findings, we predicted several potential transcripts for the M and L genes, which share the same start site S3 and terminate at either T3 or T4. The M gene could produce a short transcript encoding the M ORF, which can either remain unspliced (M t1; size: 741 nt) or undergo splicing at In1 to produce a shorter transcript (M t2; 565 nt), both theoretically terminating at T3. Although the unspliced transcript M t1 was detected by RT-PCR, it appears to be in the minority of transcripts based on RNAseq results. In addition, four transcripts were predicted, termed M-L t1-4, all sharing the same start site S3 and termination site T4, but differing in their splicing patterns. The longest unspliced transcript, M-L t1, contains the full-length M and L ORFs (5,680 nt), whereas the other transcripts (M-L t2-4) are progressively truncated by splicing at different introns, resulting in transcripts of size 5,218 nt, 5,150 nt, and 4,945 nt, respectively (**Figure 7C**). These transcripts may encode either truncated versions of the L ORF or only the M ORF.

A complementary northern blot analysis of RNA from the persistently WhSBV-infected GCSB1441 and the experimentally WhSBV-infected EPC cells, supported some findings from the sequence data analysis (**Figure 8A-C**). All probes (N, X, P, G, M, or L ORF specific) detected the full genome of WhSBV from the GCSB1441 and EPC cells at the expected size of 9,008 nt. Probes X, P and G probably labelled the same RNA species at approximately 2,100 nt, matching the predicted length for the polycistronic X-P-G mRNA. The M and L probes both labelled an RNA between 6 and 8 kb, likely representing the M-L t1 transcript. A faint band at 500 nt was detected using the M probe, but not with the L probe, which may represent the transcripts encoding the M ORF alone. The N probe identified an RNA species that was consistently higher than expected and was also recognised by the X, P and G probes. This observation suggests the presence of a RNA spanning N, X, P and G, while the monocistronic N mRNA was not detected at the expected size.

**Figure 8:**
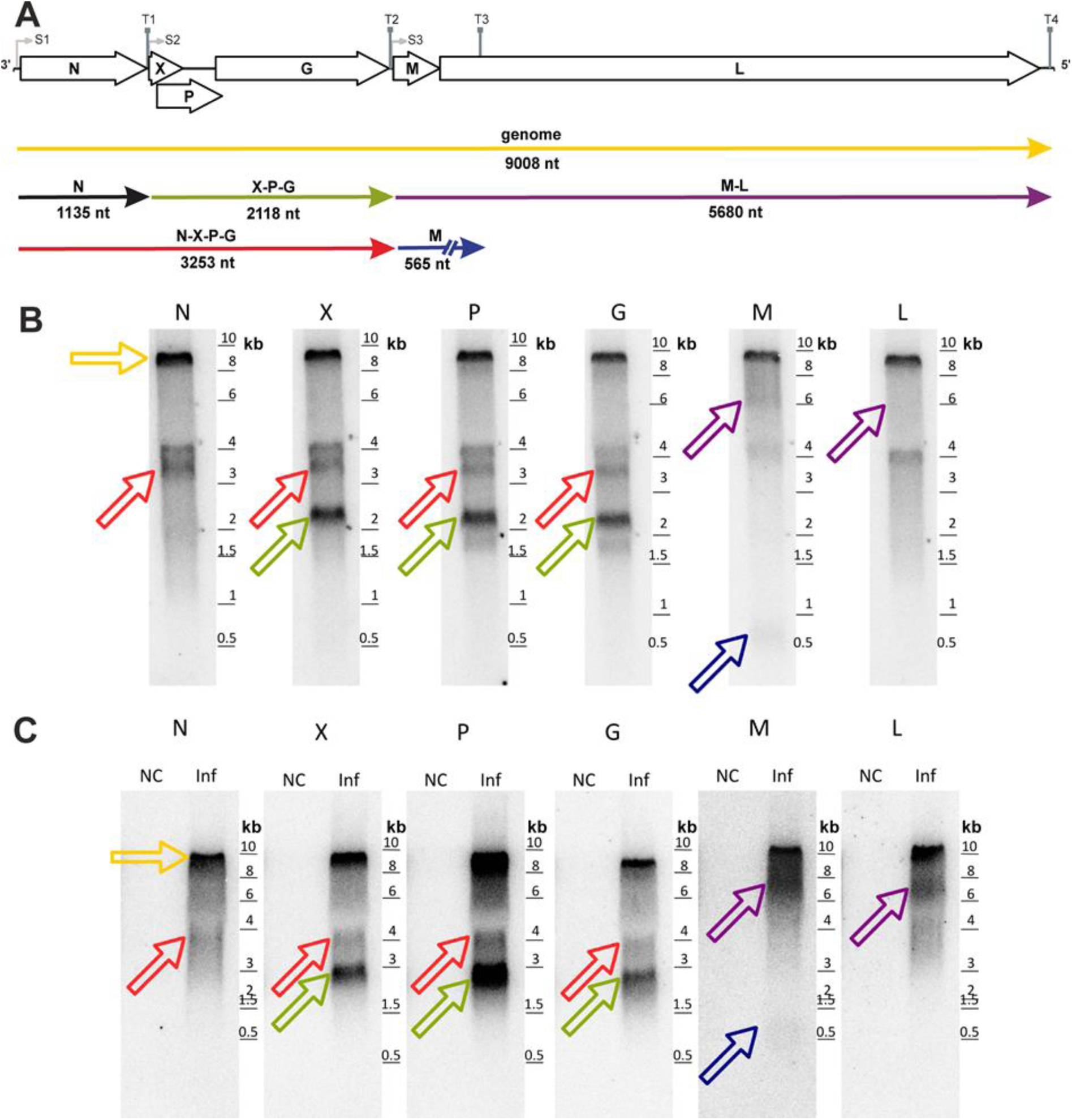
Detection of multicistronic WhSBV RNA in persistently WhSBV-infected and WhSBV-inoculated cells. WhSBV RNA transcripts were detected by Northern blot using ^32^P-CTP-labelled probes specific for WhSBV genes (arrows) encoding for the nucleoprotein (N), accessory protein (X), phosphoprotein (P), matrix protein (M), glycoprotein (G), and the large protein (L). For reference, a schematic representation of the WhSBV genome, along with open reading frames (arrows) and possible RNA molecules (coloured lines) is shown in **(A).** Total RNA from the persistently WhSBV-infected GCSB1441 cell line **(B)** and WhSBV-inoculated (Inf) EPC cells compared to non-WhSBV-inoculated (NC) EPC cells **(C)**, was used for the Northern blot. Probes that detected RNA species of the same size were interpreted as corresponding to the same RNA molecule. The coloured arrows in **(B)** and **(C)** correspond to the respective WhSBV RNA molecules in the schematic. A ladder is provided in kilobase pairs (kb).

High-resolution mass spectrometry was performed on the persistently WhSBV-infected GCSB1441 cell suspension to provide an overview of the viral proteins. The analysis revealed peptides that matched all the predicted proteins of WhSBV, except for the L ORF where no matching peptides were detected. No fused peptides were detected either, suggesting no additional splicing (**Figure 9**).

**Figure 9:**
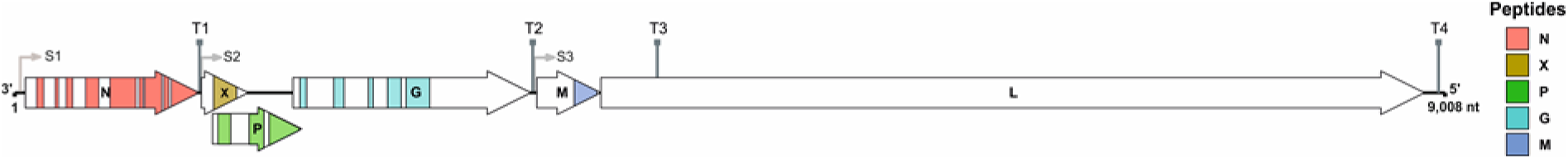
Detection of WhSBV viral peptides in persistently WhSBV-infected GCSB1441 cells. High resolution mass spectrometry was performed on protein extracts from the persistently WhSBV-infected GCSB1441 cells. Mass spectrometry raw files were processed with MaxQuant (version 2.4.13.0) [31] using a custom virus database (6 entries) and the *C. idella* database (C_idella_female_genemodels.v1; 32,811 entries) with default settings. The figure shows a schematic representation of the complete viral genome, with open reading frames (ORF) indicated by as arrows and transcription start (S1-S3) and termination sites (T1-4). Areas corresponding to detected peptides are colour coded for each associated ORF. No peptides matching the L protein or fused peptides were detected.

## DISCUSSION

This study represents the first *in vitro* molecular characterisation of Wŭhàn sharpbelly bornavirus (WhSBV), a member of the family *Bornaviridae*, genus *Cultervirus* [3, 6], and thus the first biological characterisation of a bornavirid not belonging to the well-studied genus *Orthobornavirus*. Initially, identified by RNA sequencing of sharpbelly tissues (gut, liver and gill) [2, 6], WhSBV was later detected by data mining in grass carp cell lines (CIK and L8824) [2, 6, 32]. Here, we further identified WhSBV in two grass carp swim bladder-derived cell lines by metagenomic RNA sequencing. Both cell lines were originally established by the Pearl River Fisheries Research Institute (PRFRI), Guangzhou, China. Whether the tissues used to establish the cell lines were harvested from an WhSBV-infected animal or were contaminated during handling from an unknown source remains unknown. While contamination cannot be completely ruled out, the fact that WhSBV replicates and persistently infects cypriniform cell lines suggests that the virus is more likely to have originated from the grass carp used for the cell culture. This suggests that cypriniform fish, such as grass carp and sharpbelly fish, are the preferred host of WhSBV in natural environments. However, the natural host range of WhSBV and other culterviruses remains unknown.

The identification of WhSBV in established cell lines allowed us to study the virus *in vitro*, offering insights into WhSBV persistence, potential *in vitro* host range, and transcriptional strategies. Although the lack of WhSBV-specific antibodies and the absence of a cytopathic effect in cell culture limited the ability to conduct classic virus titration, we used RT-qPCR and RNA ISH instead. Other studies have shown good correlation between viral titres and Cq values, enabling a semi-quantitative assessment of viral load to be made in the absence of virus titration [33–35]. Our RT-qPCR assay targeting the G gene was the most reliable and sensitive method of detecting WhSBV RNA. This is consistent with the findings of Northern blot analysis and genome read coverage, which showed that the X-P-G coding RNA was expressed at higher levels than other viral RNAs.

Our comparative analysis across cypriniform cell lines revealed that WhSBV can establish a persistent infection without inducing cytopathic effects and maintains stable viral RNA levels over multiple passages (**Figure 2A**). Similar behaviour has been observed for BoDV-1, BoDV-2, VSBV-1 and several avian orthobornaviruses, where some inoculated cell lines became persistently infected without cytopathic effect and could be sub-cultivated without loss of infectivity [36–43]. As some orthobornaviruses show relatively broad host spectrum *in vitro* we tested WhSBV inoculation of non-cypriniform, reptile, avian and mammalian cell lines.

Avian and mammalian cells were clearly not susceptible to WhSBV infection. This may be due either to the absence of essential host factors in avian and mammalian cells required for WhSBV replication, the activation of an effective antiviral response, or the virus’s potential intolerance to the elevated incubation temperature of 37°C.

In some non-cypriniform fish (RTG-2/f, CHSE-214, and TiB) and reptilian cell lines, susceptibility to WhSBV infection was difficult to assess because ACTB levels declined over several passages, with eventual loss of cultures. This phenomenon was also observed in uninfected control cultures, suggesting the deterioration was likely due to suboptimal culture conditions, rather than WhSBV infection. Nevertheless, in these cell lines, viral RNA levels over the successive passages fluctuated near or below the detection Cq cut-off value. While these low-level signals may reflect residual inoculum, RNA ISH confirmed the presence of viral RNA within some cells for RTG-2/f and TiB, indicating that these cells were indeed infected by WhSBV. However, infection remained confined to individual cells and there was no progressive increase in the percentage of infected cells, unlike in cypriniform cells where infection spread over time. As RNA ISH was not possible for reptilian cells, there is currently no conclusive evidence whether these are susceptible to WhSBV infection and further investigations will be necessary.

The difference in WhSBV infectivity between cypriniform and non-cypriniform fish cell lines may be attributed, in part, to inherent cytological and physiological distinctions between these cells. The fish cell lines used in this study originate from various tissues and primarily represents epithelial or fibroblast lineages. These cells may differ substantially in membrane composition, receptor expression profiles, and basal innate immune activity, all of which can influence their permissiveness to viral entry, replication, and persistence [44]. Furthermore, variation in metabolic activity, temperature tolerance, and stress responses can further modulate viral propagation *in vitro* [45–48]. Host-range restrictions of this kind are not unique to WhSBV. For instance, infectious hematopoietic necrosis virus (IHNV; family *Rhabdoviridae*) exhibits strong host specificity and replicates robustly in salmonid-derived cell lines, such as, RTG-2 (rainbow trout; gonad) and CHSE-214 (Chinook salmon; embryo), but shows limited susceptibility in non-salmonids like EPC (fathead minnow; *Epithelioma papulosum*) and FHM (fathead minnow; muscle) [49–52]. Similarly, cyprinid herpesvirus 3 (CyHV-3; family *Alloherpesviridae*) demonstrated pronounced tropism toward cyprinid hosts and replicates efficiently in certain cypriniform cell lines derived from common carp and koi but fails to establish productive infection in the cypriniform EPC cells and non-cypriniform cell lines as those from catfish [44]. These examples underscore the critical role of taxonomic relatedness and cellular context in shaping viral host range and tissue tropism. Further studies are warranted to elucidate the precise mechanisms underlying WhSBV’s restricted replication in these non-cypriniform cell types.

The restricted replication of WhSBV in non-cypriniform fish cells may indicate an evolutionary history of adaptation to cypriniform hosts, and therefore the potential existence of other related culterviruses with a greater affinity for non-cypriniform hosts. An *in silico* screening identified five novel fish bornavirids in bony and cartilaginous fish hosts that showed significant genomic divergence, suggesting that the diversity of culterviruses is currently underestimated [2]. Nevertheless, current *in silico* and metagenomics evidence indicates that both WhSBV and Murray-Darling carp bornavirus (MDCBV) are restricted to cypriniform hosts, with no confirmed detections in other taxonomic groups to date [2, 6, 32]. These findings suggest a potential evolutionary adaption of WhSBV and MDCBV to cypriniform species. Future studies will be essential to determine host range of WhSBV and other culterviruses in natural settings.

For production of virus preparations, active disruption of the infected cells by e.g. freeze/thawing was necessary, while comparably little infectious virus appeared to be present in the supernatant of infected culture. This is in agreement with orthobornaviruses, such as BoDV-1 [53, 54]. We observed virus particle-like structures and budding in WhSBV-inoculated ZF4 cells, which were absent in the respective controls. Although these structures cannot be definitively attributed to WhSBV, their morphology is consistent with that of viral particles. With 150-180 nm in diameter, their detected size exceeds the typical size range reported for BoDV-1 (70-130 nm) [55, 56] and avian bornaviruses (60-104 nm) [57–59]. While the observed structures may represent viral particles and budding, it is currently not possible to confirm their identity as WhSBV without specific staining tools, such as immunogold labelling.

The innate antiviral immune system, including the interferon (IFN) pathway, is well conserved between mammals and teleost fish [60–62]. In teleost, as in higher vertebrates, viral nucleic acids are sensed by pattern recognition receptors (PRRs), notably RIG-I-like receptors such as RIG-I and MDA5 [63–65]. Upon sensing, these receptors initiate signalling cascades that culminate in the production of type I IFNs and pro-inflammatory cytokines, which then upregulate interferon-stimulated genes (ISGs), to restrict viral replication and enhance viral clearance [66, 65, 67–69]. In order to investigate the cellular response to WhSBV infection and to gain insight into the possible mechanisms underlying its ability to establish a persistent infection without inducing cytopathic effects, we analysed the host gene expression at different time points of infection. Transcriptomic analysis of the experimentally WhSBV-infected CCB cells indicated a limited initial activation of the innate immune response shortly after inoculation at 4 and 8 hpi by upregulation of the genes IRF1B and RASAL2 (**Figure 6E**). Interferon regulatory factors, such as IRF1B, are key regulators of the innate antiviral response in vertebrates, controlling the expression of type I IFNs and ISGs [70, 71]. In zebrafish, IRF1B alone was shown to induce the expression of IFNs and ISGs, resulting in effective protection against viral infection [72]. While the role of RASAL2 in fish remains unclear, though in mammals, it has been characterized as tumour and metastasis suppressor [73]. Despite this initial immune activation, no significant changes in host cell gene expression were observed between 24 and 96 hpi (**Figure 6C**), even as viral RNA levels and the proportion of infected cells increased steadily during this period (**Figure 6B**). These findings suggest that WhSBV may evade or suppress early innate immune signalling through yet-unknow mechanisms, thereby enabling efficient viral replication and the establishment of persistent infection. In comparison, orthobornaviruses have been shown to evade IFN induction by possessing monophosphorylated genome ends [74, 75]. BoDV-1 further suppresses the interferon response via P-protein-mediated inhibition of TANK-binding kinase 1 (TBK-1) [76] and N-protein interference with IRF7 activation [77].

Only at 120 and 240 hpi we observed the upregulation of RIG-I and IRFs other than IRF1B, that may have contributed to a delayed innate immune response at later stages. This response was marked by significant upregulation of many ISGs, such as MxB, GVINP1, ISG15, and RSDA2, all of which play antiviral roles in teleost fish [78]. This suggests that the host’s innate immune system did not mount a response until the infection had spread to nearly all cells. However, this late response appeared insufficient to clear the infection, as the cells remained persistently infected, suggesting either the activation of a secondary immune response aimed to control viral load, or that the increasing intracellular viral burden eventually triggered the innate immune system of the host. Despite the upregulation of ISGs, no significant induction of apoptosis-or necrosis-related genes was observed throughout infection, suggesting that WhSBV establishes infection without inducing programmed cell death. This may contribute to continued viral replication and long-term or even persistent infection. In contrast, common carp hepatocytes exposed to ammonia stress undergo apoptosis via the P53-BAC/BCL-2 pathway, demonstrating the species’ ability to induce cell death under certain conditions [79]. The absence of such responses in WhSBV-infected cells raised the possibility that the virus actively suppresses apoptosis. Our study provides a temporal overview of host gene expression during a 10-day infection course in the cypriniform CCB cells. To fully elucidate the molecular strategies employed by WhSBV to evade and/or modulate the host immune response, future investigations should focus on identifying the specific viral genes, proteins or genomic structures involved in immune suppression and persistence. The establishment of persistence without cytopathic effects is a common feature of orthobornaviruses [3] and has also been observed in other mononegaviruses, such as arenavirids (family *Arenaviridae*), which suppress host antiviral responses, particularly by inhibiting type I interferon production, to maintain chronic infections [80, 81].

One strategy to evade the hosts immune response might be the strict regulation of WhSBVs transcription, that involves expression gradients, polycistronic mRNAs and splicing, as seen in some orthobornaviruses [82, 83, 11, 12]. Some regulatory features, such as the splicing of M-L transcripts, and the position and sequence of transcription start and termination sites [54, 11], seem to be highly conserved among culterviruses and therefore suggest conserved mechanisms. This transcription strategy is in contrast to the predominantly monocistronic transcription seen in other mononegaviruses [84], such as paramyxovirids (family *Paramyxoviridae*), where the viral genome is transcribed into monocistronic unspliced mRNAs, each encoding a single protein [85]. However, there are some exceptions such as Ebola virus (family *Filoviridae*), that uses RNA editing of its glycoprotein gene in order to generate multiple variants, thereby optimising replication and pathogenicity without increasing genome size [86].

Northern blot and sequence coverage analysis indicated that the N proteins are expressed not only from a monocistronic mRNA, but also co-expressed with X/P and G from a polycistronic mRNA. However, it remains to be clarified whether the detected RNA of approximately 3000-4000 nt in length is actually mRNA or a sub-genomic RNA. For BoDV-1, the non-coding region between the N and X genes shows structural similarities to that of WhSBV, where T1 is located downstream of the transcription start site S2 in BoDV-1 [87]. The consensus motif of the BoDV-1 termination sites, U(A)_6-7_ [88], serves as a core signal sequence but is insufficient for efficient transcription termination on its own, as the surrounding upstream and downstream nucleotides significantly influence T1 utilisation. Consequently, when transcription is initiated at S1 and terminated at T1, these regulatory elements contribute to the production of a monocistronic (1,200 nt) mRNA encoding the N protein [87]. However, studies have demonstrated that the viral RNA-directed RNA polymerase frequently bypasses the T1 transcription termination site, resulting in the production of a longer (1,900 nt) polycistronic mRNA containing the N, X, and P ORFs [87]. The presence of such transcripts in BoDV-1-infected cell cultures and rat brains suggests that they are likely authentic mRNAs, rather than by-products of aberrant replication, and their production may be related to viral fitness [87]. Conversely, other studies have suggested that these long mRNAs may represent aborted replication products or an extended form of “leader RNA”, similar to those described in certain other negative-strand RNA viruses [10, 89]. Therefore, future studies are needed to investigate the function of these multicistronic mRNAs, particularly those co-expressing N with X/P and G, and to determine whether they encode proteins or contribute to viral fitness.

The relatively low expression level of the L gene observed in WhSBV may be explained by a combination of transcriptional and post-transcriptional mechanisms. One potential factor is the presence of a transcription termination signal (T3) located within the L ORF. In BoDV-1, the T3 signal mediates premature transcription termination of the L ORF, resulting in lower transcript abundance and consequently lower protein expression [10]. A similar regulatory structure in WhSBV could result in early termination of L gene transcripts, as supported by our transcriptional profiling, which show high coverage starting at S3 and ending at T3. Additionally, our PCR and Sanger sequencing analyses revealed the presences of introns within the L ORF, suggesting splicing event that may disrupt the integrity of the L ORF. This could serve to downregulate L expression while allowing expression of the overlapping M ORF. This phenomenon has also been reported in BoDV-1, where alternative splicing within the L gene results in expression of the overlapping G ORF [82, 8, 11]. Although L transcripts were detected by Northern blot, proteomic analysis did not identify the L protein, further supporting its comparable low expression levels. Collectively, these findings suggest that the L gene in WhSBV may be tightly controlled to fine-tune L protein levels. Future studies using more sensitive proteomic techniques and functional assays are warranted to clarify whether L protein expression is truly low or simply undetectable under current conditions. Additionally, further investigation is needed to elucidate the functional relevance of the observed introns. Based on the current data, possible hypotheses include: facilitating increased expression of M gene by truncating the L ORF, regulating L expression at low levels, or generating an alternative transcript encoding a yet-unidentified protein. A similar mechanism has been observed for vesicular stomatitis virus (VSV; family *Rhabdoviridae*), which controls L gene expression by polycistronic transcription, co-expressing it with other viral genes to balance protein levels and minimise host cell stress while optimising viral replication and assembly [90]. This specific transcriptional regulation of WhSBV may balance viral protein levels but requires further investigation.

## CONCLUSION

This study provides a molecular characterisation of WhSBV, highlighting its evolutionary adaptation to cypriniform hosts. The virus replicates efficiently in cypriniform cell cultures, establishing persistent infections without inducing any visible cell damage and suppressing the host’s early innate immune responses. Its limited or absent replication in non-cypriniform cultures highlights the need for *in vivo* validation to determine its full host range. Additionally, this study sheds light on the transcriptional mechanisms of WhSBV, revealing its regulatory strategies for modulating gene expression. Further research is essential to unravel the molecular mechanisms underlying WhSBV persistence and to assess its ecological impact, particularly in aquaculture, where persistent viral infections could significantly impact fish health and disease management. Our findings also emphasise the importance of transitioning from *in silico* screening to *in vitro* validation — a critical step in bioinformatics-driven virology research. Although metagenomic approaches facilitate virus discovery, a key limitation is often the lack of direct validation from real virus samples.

## DATA AVAILABILITY

All sequencing data associated with this study have been deposited in public repositories and are freely accessible. The WhSBV genome sequence is available in GenBank under the accession number PV171101. Additional metadata can be found under BioProject PRJNA1226401 and BioSample SAMN46928747. Raw sequencing reads of WhSBV persistently infected GCSB1441 are available in the Sequence Read Archive (SRA) under SRR32424572, and RNAseq data have been deposited at ArrayExpress under E-MTAB-14894.

## ACKNOWLEGEMENTS

The investigations were supported by a Friedrich-Loeffler-Institut internal PhD program, grant number FLI-IVD-XX-2021-83, granted to F.P.

We gratefully acknowledge the assistance of Fermin Georgio Lorenzen-Schmidt, Mathias Lenk from the Collection of Cell Lines in Veterinary Medicine (CCLV), and the contributions of Leoni Lemm, Robin Brandt, and Kristin Vorpahl.

## CONFLICT OF INTEREST

The authors declare no conflicts of interest. The funders had no role in study design, data collection, analysis, interpretation, manuscript writing, or decision to publish the findings.

## Notes

### Competing Interest Statement

The authors have declared no competing interest.

### Summary of Updates

We revised the manuscript following reviewer suggestions.

