## Supplementary Material for "From *in silico* to *in vitro*: Wŭhàn sharpbelly bornavirus infects and persists in cyprinoform cells"

**for**

Eshak *et al.* 2025

### Supplementary Figures

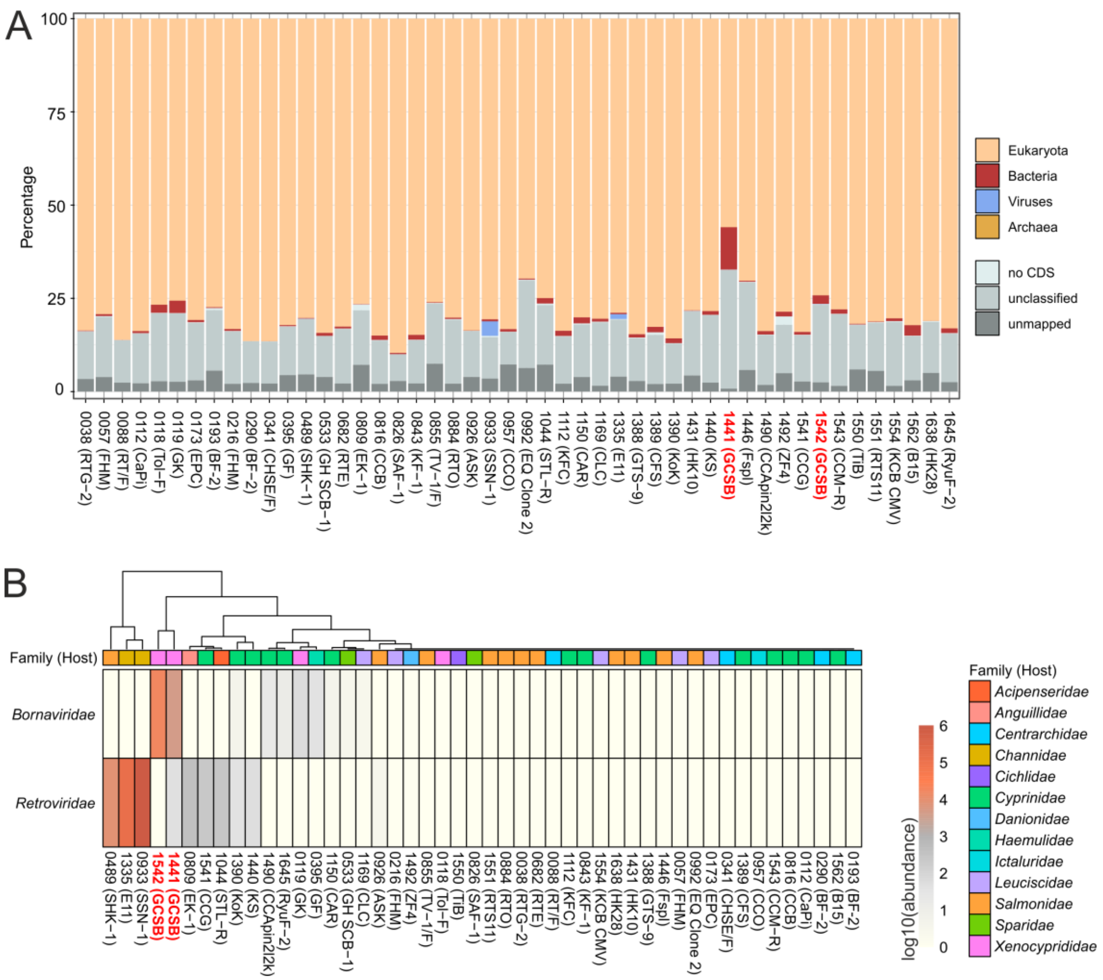


**Supplementary Figure S1**: **Metagenomic analysis of different fish cell lines.** (**A**) Using RNA sequencing and taxonomic assignment, the sequencing reads of each cell culture were grouped into the top levels of the taxonomy database maintained by NCBI/GenBank: Archaea, Eukaryota, Bacteria and Viruses. (**B**) The normalised read abundances for the taxon Viruses showed high levels of *Retroviridae* in some cell cultures. Reads of the family *Bornaviridae* were found only in the GCSB1441 and GCSB1542 cultures (highlighted in red).

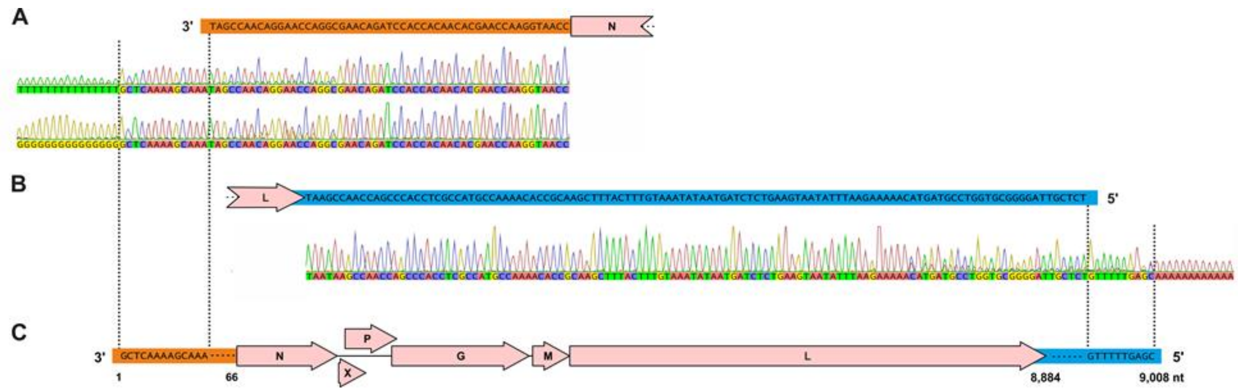


**Supplementary Figure S2**: **Confirmation of viral genome termini.** The initial WhSBV genome termini resulting from *de novo* assembly were confirmed as correct and further extended using 5'-(**A**) and 3'-RACE (**B**). The final WhSBV genome had a size of 9,008 nt (**C**), with the 5' untranslated region (UTR) and 3'-UTR being 66 and 125 nt in size, respectively.

| Cell line | Template | Assay type | N | ACTB | P | ACTB | G | ACTB | M | ACTB | L | ACTB |
|---|---|---|---|---|---|---|---|---|---|---|---|---|
| GK (0119) | RNA / DNA | qPCR (only DNA) | n.d. | 27.5 | n.d. | 26.8 | n.d. | 26.8 | n.d. | 27.0 | n.d. | 26.6 |
| GCSB1542 | RNA / DNA | | n.d. | 30.4 | n.d. | 30.5 | n.d. | 30.9 | n.d. | 30.2 | n.d. | 29.5 |
| GCSB1441 | RNA / DNA | | n.d. | 33.1 | n.d. | 32.4 | n.d. | 31.4 | n.d. | 32.3 | n.d. | 31.0 |
| GCSB1441 | cDNA | | 21.1 | n.d. | 21.2 | n.d. | 19.1 | n.d. | 20.3 | n.d. | 21.8 | n.d. |
| GK (0119) | RNA / DNA | RT-qPCR (RNA+DNA) | n.d. | 24.4 | n.d. | 24.7 | n.d. | 24.7 | n.d. | 24.3 | n.d. | 24.6 |
| GCSB1542 | RNA / DNA | | 21.5 | 24.7 | 22.0 | 26.0 | 21.2 | 24.6 | 21.6 | 26.8 | 21.6 | 25.7 |
| GCSB1441 | RNA / DNA | | 23.0 | 26.1 | 23.3 | 33.1 | 22.4 | 26.2 | 23.1 | 28.5 | 23.1 | 27.2 |

**Supplementary Figure S3: WhSBV is not integrated into the grass carp genome.** To exclude that the detected WhSBV sequences originated from endogenous viral elements (EVE) in the genomes of grass carp cells, a mixture of extracted DNA and RNA was tested using different PCR assays. Grass carp cells GCSB1441 and GCSB1542 were considered to be persistently infected with WhSBV, while grass carp cells GK were used as a negative control. In detail, primers and probes detecting the viral genes N, P, G, M, L and the cellular *ACTB* gene were used in a quantitative PCR (qPCR) without a reverse transcription (RT) step, thus amplifying only the DNA present, but not the RNA. Specific WhSBV cDNA was used as a positive control. The samples were re-tested in a PCR with an RT step (RT-qPCR) using the same prime/probe mixes. Results are presented as quantification cycles (Cq). n.d.: not detected.

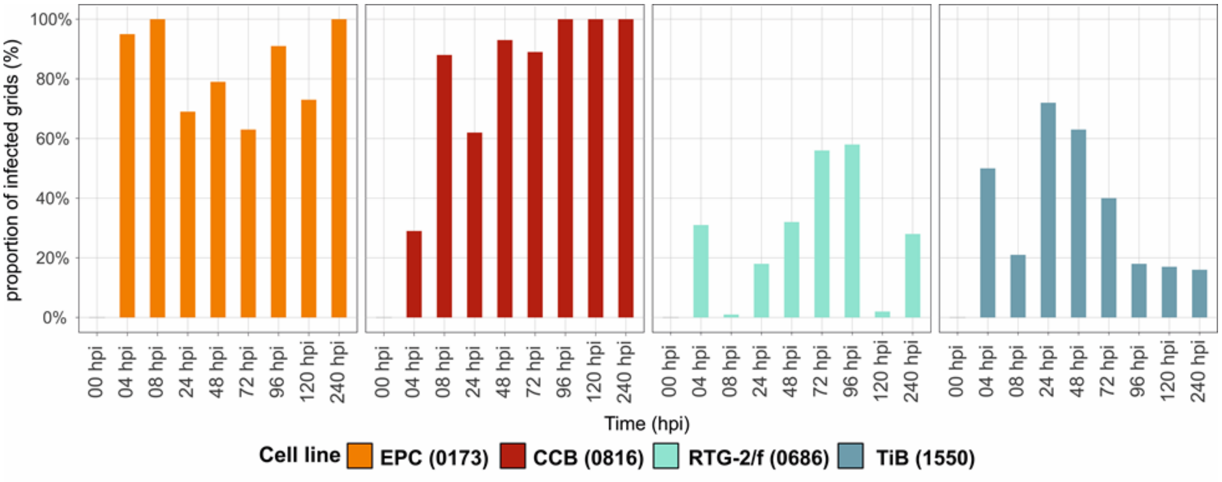


**Supplementary Figure S4: Proportion of WhSBV infected cells using RNA ISH.** Cypriniform (EPC, CCB) and non-cypriniform (RTG-2/f, TiB) cell lines were infected with WhSBV at a 1:1.7 ratio, harvested at time points between 0 and 240 hours post infection (hpi), and fixed in 10% neutral buffered formalin. Paraffin-embedded 4-µm sections were processed for RNA *in situ* hybridisation (RNA ISH) using RNAScope with WhSBV-specific probes, alongside technical controls. The proportion of infected cells was quantified as a percentage in a 10 × 10 grid (size 100 × 100 µm per grid cell).

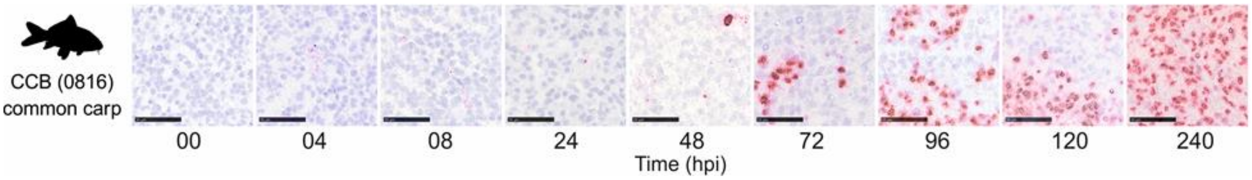


**Supplementary Figure S5: Detection of viral (+)RNA by RNA *in situ* hybridisation.** RNA *in situ* hybridisation using a custom-designed probe against WhSBV (+)RNA (anti-genome, mRNA) was performed on formalin-fixed cell pellets from the WhSBV-inoculated cypriniform cell line CCB harvested at time points between 0 and 240 hours post infection (hpi). A representative image is shown for each time point. The black scale bar represents 50 µm. Early infection was characterised by fine-granular cytoplasmic labelling, whereas larger, globular signals appeared at later time points. During the course of the experiment, single positive cells were observed initially, followed by foci of positive cells, which eventually became confluent and resulted in diffuse labelling of the cell pellet.

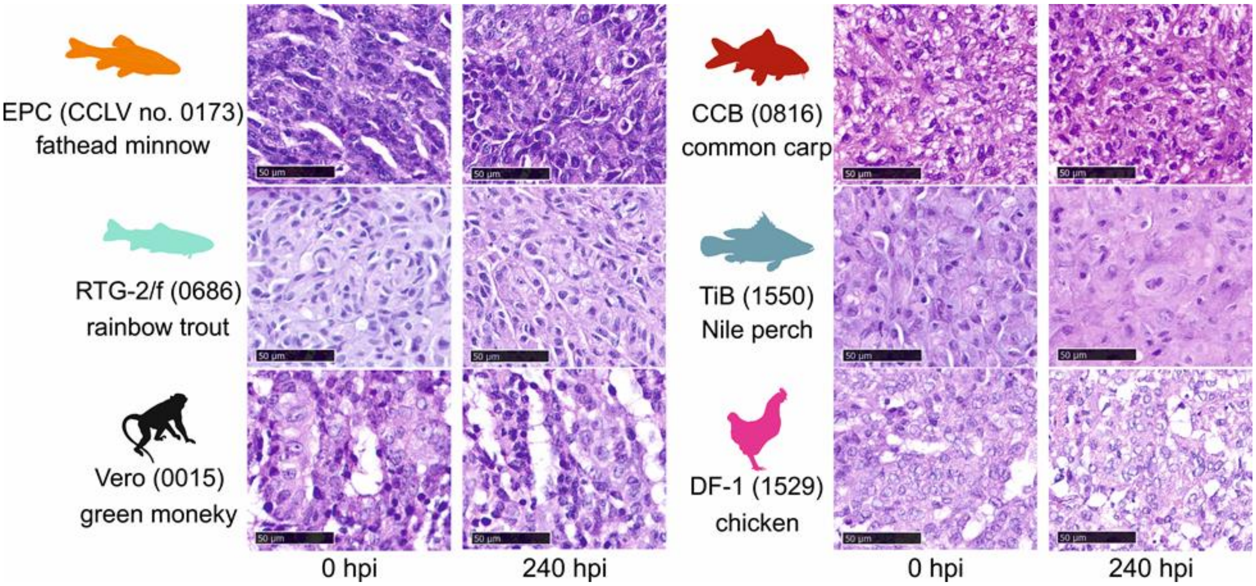


**Supplementary Figure S6: Histopathology** of formalin-fixed, paraffin-embedded and hematoxylin-eosin stained cell pellets from WhSBV-inoculated cell lines. The black scale bar represents 50 µm.

### Supplementary Tables

**Supplementary Table S1:** Summary of cell lines used for metagenomic RNA sequencing

**Supplementary Table S2:** Primers and probes used in this study

**Supplementary Table S3:** Complete list of cell lines used for infection experiments in this study

**Supplementary Table S4:** Summary of differentially expressed genes

**Supplementary Table S5:** Transcription start and termination motifs of WhSBV
